# Central amygdala circuits modulate food consumption through a positive valence mechanism

**DOI:** 10.1101/145375

**Authors:** Amelia M. Douglass, Hakan Kucukdereli, Marion Ponserre, Milica Markovic, Jan Gründemann, Cornelia Strobel, Pilar L. Alcala Morales, Karl-Klaus Conzelmann, Andreas Lüthi, Rüdiger Klein

## Abstract

The complex behaviors underlying the pursuit and consumption of rewards are integral to an organism’s survival. The hypothalamus and mesolimbic dopamine system are key mediators of these behaviors, yet regulation of appetitive and consummatory behaviors outside of these regions is not well understood. The central nucleus of the amygdala (CeA) is implicated in feeding and reward behavior, but the specific neural players and circuit mechanisms that positively regulate these behaviors remain unclear. Here, we define the neuronal mechanisms by which the CeA promotes consumption of food. We show, using *in vivo* activity manipulations and Ca^2+^ imaging, that CeA GABAergic neurons expressing the serotonin receptor 2a (Htr2a) modulate food consumption in multiple contexts, promote positive reinforcement and are active *in vivo* during eating. We demonstrate using electrophysiology, anatomical tracing methods and *in vivo* optogenetics that both intra-CeA and long-range circuit mechanisms underlie these functional effects. Finally, we show that CeA^Htr2a^ neurons are poised to regulate food consumption through inputs from feeding-relevant brain regions. Our study highlights a mechanism by which defined CeA neural circuits positively regulate food consumption.

## Introduction

Survival of an organism relies on the ability to seek out and consume natural rewards in an ever-changing environment. To do so, animals must attend to salient environmental stimuli and execute complex motivated behavioral sequences to successfully seek and consume the reward. A network of brain regions, involving most notably the lateral hypothalamus (LH), ventral tegmental area (VTA) and nucleus accumbens (NAc), mediate goal-directed reward seeking and consumption behavior by increasing the saliency of environment cues and promoting positive reinforcement^1–4^. The CeA, a forebrain structure of striatal origin, is important for processing salient stimuli and orchestrating the appropriate behavioral response. This region is comprised of a highly interconnected network of inhibitory GABAergic neurons that are functionally classified based on expression of molecular markers, several of which have described roles in fear ^5–8^, anxiety ^9,10^ and appetite suppression ^11^. Although the CeA has a reported role in magnifying reward saliency^12–14^, modulating food consumption^11,14^, and promoting appetitive behaviors^15^, the cellular heterogeneity and high degree of neural interconnectivity in this region has precluded an insight into the specific neural players and underlying circuits that positively regulate food consumption.

Here we report that a molecularly defined population of CeA neurons positively modulates food consumption. Using optogenetic and pharmacogenetic tools, we demonstrate that CeA neurons expressing the serotonin receptor Htr2a promote food consumption and positive reinforcement. Deep brain calcium imaging revealed that CeA^Htr2a^ neurons consistently increase activity during eating. We show, using optogenetic and rabies tracing techniques, a local intra-CeA circuit mechanism by which CeA^Htr2a^ neurons exert their functions. Further, we reveal that CeA^Htr2a^ neurons promote feeding and positive reinforcement through long-range inhibition of cells within the parabrachial nucleus (PBN), a brain region known to process aversive gustatory and sensory signals ^16–18^. Finally, we demonstrate that PBN-projecting CeA^Htr2a^ neurons receive distinct monosynaptic inputs from brain regions with known roles in feeding behaviors. Together these findings reveal the specific neural players within the CeA that positively regulate eating behavior.

## Results

### CeA^Htr2a^ neurons modulate food consumption

To identify CeA neural subpopulations that positively regulate food consumption, we searched the GENSAT transgenic mouse collection for a mouse line that permitted genetic access to CeA neurons mutually exclusive of the anorexigenic population marked by expression of protein kinase C-δ (PKCδ)^11^. Of the top candidates, the BAC transgenic line *Htr2a-Cre* (KM208) was found to be highly expressed within the CeA. Our characterization of the line by breeding *Htr2a-Cre* mice to a LacZ reporter line confimed the faithful representation of Htr2a+ neurons in the CeA (Supplementary Fig. 1a, b) and revealed no overlap with CeA^PKCδ^ neurons (Fig. 1a, b), but partial overlap with other CeA genetic markers (Supplementary Fig. 1c-f). Physiologically, CeA^Htr2a^ neurons were found to be homogeneous and exhibited late firing properties (Fig. 1c, d). Given that CeA^Htr2a^ and CeA^PKCδ^ neurons were mutually exclusive populations, we determined whether CeA^Htr2a^ neurons promoted food intake. First, we virally targeted CeA^Htr2a^ neurons in *Htr2a-Cre* mice with Cre-dependent stimulatory hM3Dq designer receptors exclusively activated by designer drugs (DREADDs)^19,20^ and performed a free-feeding assay (Fig. 1e, f and Supplementary Fig. 2a-d). Acute activation of CeA^Htr2a^ neurons by intraperitoneal injection of the DREADD ligand clozapine-N-oxide (CNO) in satiated mice increased food intake compared to controls by increasing the total time that the animals spent feeding (Fig. 1g and Supplementary Fig. 2e-h). The animals preferred food over clay pellets of similar size and hardness to the food (Fig. 2h and Supplementary Fig. 2i-k), indicating that activation of CeA^Htr2a^ neurons specifically led to food intake rather than ill-directed consummatory behavior.

**Figure 1.**
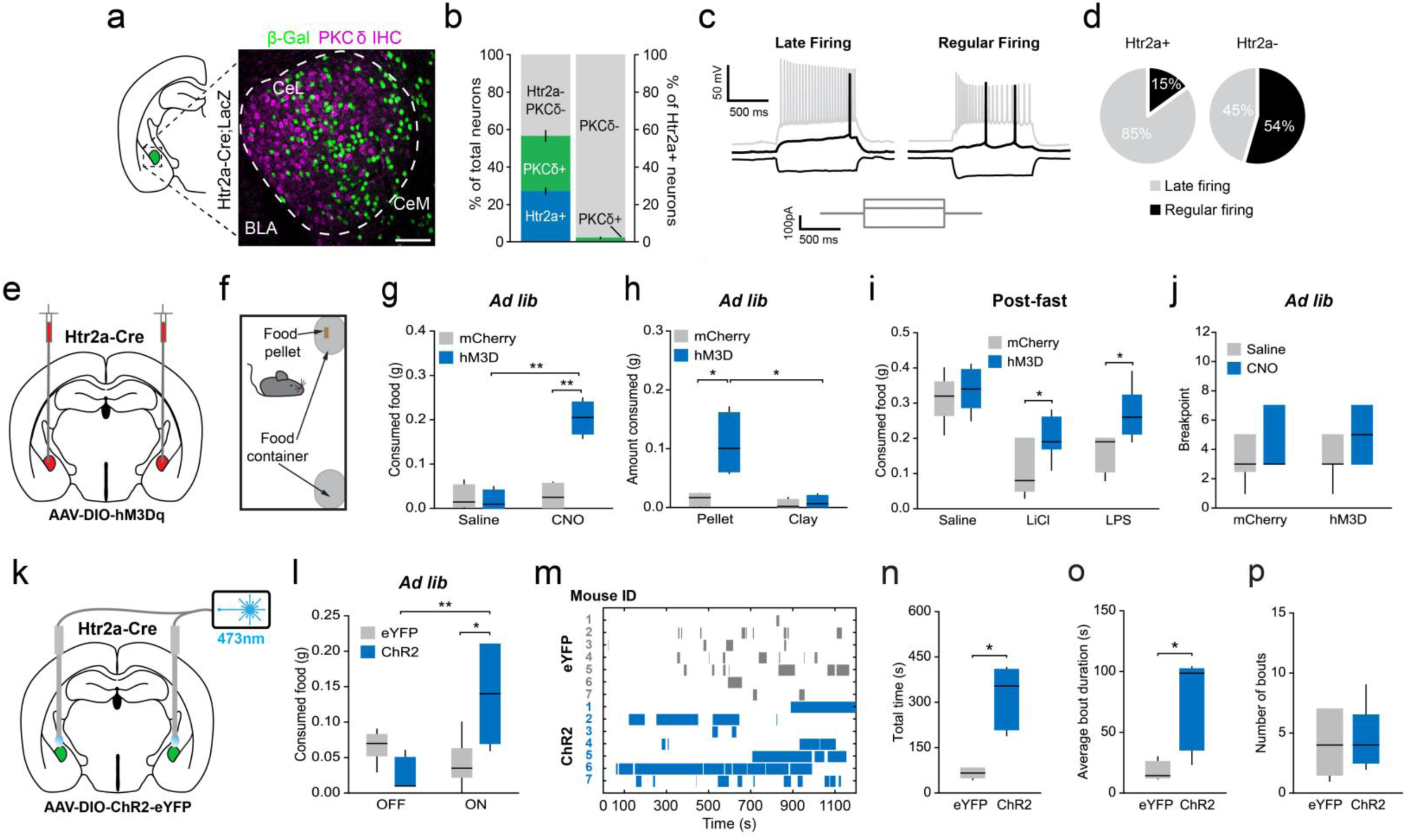
CeA^Htr2a^ neurons increase food consumption. **a**, Representative image of the CeA from *Htr2a-Cre;floxed-lacZ* mouse immunostained for-β-Gal and PKCδ. CeL, central lateral amygdala, CeM, central medial amygdala; BLA, basolateral amygdala. **b**, Quantification of the percentage of CeA neurons marked by Htr2a and PKCδ (left) and the percentage of CeA^Htr2a^ neurons that express PKCδ (right) (n = 3 mice). **c**, Whole-cell current clamp recordings from CeA^Htr2a+^ and CeA^Htr2a^‒ neurons. Lower: current injection steps. Left: example voltage trace of late firing neuron. Right: example trace of regular firing neuron. **d**, The majority (16/20) of CeA^Htr2a+^ neurons are late firing, whereas CeA^Htr2a^‒ neurons are comprised of both late (5/11) and regular firing neurons (6/11). **e**, Viral delivery of AAV hM3Dq-mCherry into the CeA of *Htr2a-Cre* mice. **f**, Scheme of free-feeding assay. **g**, Chemogenetic activation of CeA^Htr2a^ neurons increased food consumption in *ad libitum* fed mice (n = 6, two-way ANOVA: Virus: F(1,20) = 11.62, p = 0.0028, Drug: F(1,20) = 5.75, p = 0.0263, Interaction: F(1,20) = 5.84, p = 0.0253, Bonferroni post-hoc analysis: **p<0.01). **h**, Chemogenetic activation of CeA^Htr2a^ neurons increased consumption of food compared to a clay pellet (n = 6, two-way ANOVA: Virus: F(1,20) = 5.06, p = 0.0357, Food: F(1,20) = 4.72, p = 0.0419, Interaction: F(1,20) = 3.02, p = 0.0978, Bonferroni post-hoc analysis: *p<0.05). **i**, Chemogenetic activation of CeA^Htr2a^ neurons increased food intake in food-deprived mice after lithium chloride (LiCl) (n = 7 mCherry (Saline), n = 8 hM3D (Saline), n = 9 mCherry (LiCl), n = 11 hM3D (LiCl), two-way ANOVA: Virus: F(1,31) = 3.60, p = 0.0670, Drug: F(1,31) = 50.77, p = <0.0001, Interaction: F(1,31) = 2.41, p = 0.1310, Bonferroni post-hoc analysis: *p<0.05) and lipopolysaccharide (LPS) injection (n = 7 mCherry (Saline), n = 8 hM3D (Saline), n = 8 mCherry (LPS), n = 8 hM3D (LPS), two-way ANOVA: Virus: F(1,26) = 4.76, p = 0.0384, Drug: F(1,26) = 9.96, p = 0.0040, Interaction: F(1,26) = 3.37, p = 0.0780, Bonferroni post-hoc analysis: *p<0.05). **j**, Chemogenetic activation of CeA^Htr2a^ neurons did not increase effort to obtain food rewards in a progressive ratio 2 task (n= 8 mCherry, n= 9 hM3D, two-tailed paired t-test: mCherry; t(7) = 0.81, p = 0.4423, hM3D; t(8) = 0.97, p = 0.3584). **k**, Optic fiber placement above CeA^Htr2a^ ::ChR2-expressing neurons. **l**, Quantity of food consumed by *ad libitum* fed CeA^Htr2a^::ChR2 and control mice during 20 Hz photostimulation and non-photostimulated 20 minute epochs (n = 8 eYFP, n = 9 ChR2, eYFP vs ChR2 ON, two-tailed unpaired t-test: t(15) = 2.87, p = 0.0118; ChR2 OFF vs ON, two-tailed paired t-test: t(15) = 3.45, p = 0.0087). **m**, Raster plot of feeding bouts of example individual mice. **n**, CeA^Htr2a^::ChR2 mice subjected to 20 Hz photostimulation spent more time feeding (n = 6 eYFP, n = 5 ChR2, two-tailed unpaired t-test: t(12) = 2.72, p = 0.0186). **o**, The average duration of the feeding bouts was increased in CeA^Htr2a^::ChR2 mice (n = 6 eYFP, n = 5 ChR2, Mann Whitney test: p = 0.0175). **p**, The number of feeding bouts was not significantly increased in CeA^Htr2a^::ChR2 animals compared to CeA^Htr2a^::eYFP controls feeding (n = 6 eYFP, n = 5 ChR2, two-tailed unpaired t-test: t(12) = 0.23, p = 0.8224). Box–whisker plots display median, interquartile range and 5th–95th percentiles of the distribution. Bar graphs indicate mean ± SEM. *p<0.05, **p<0.01. Scale bar: 100µm.

**Figure 2.**
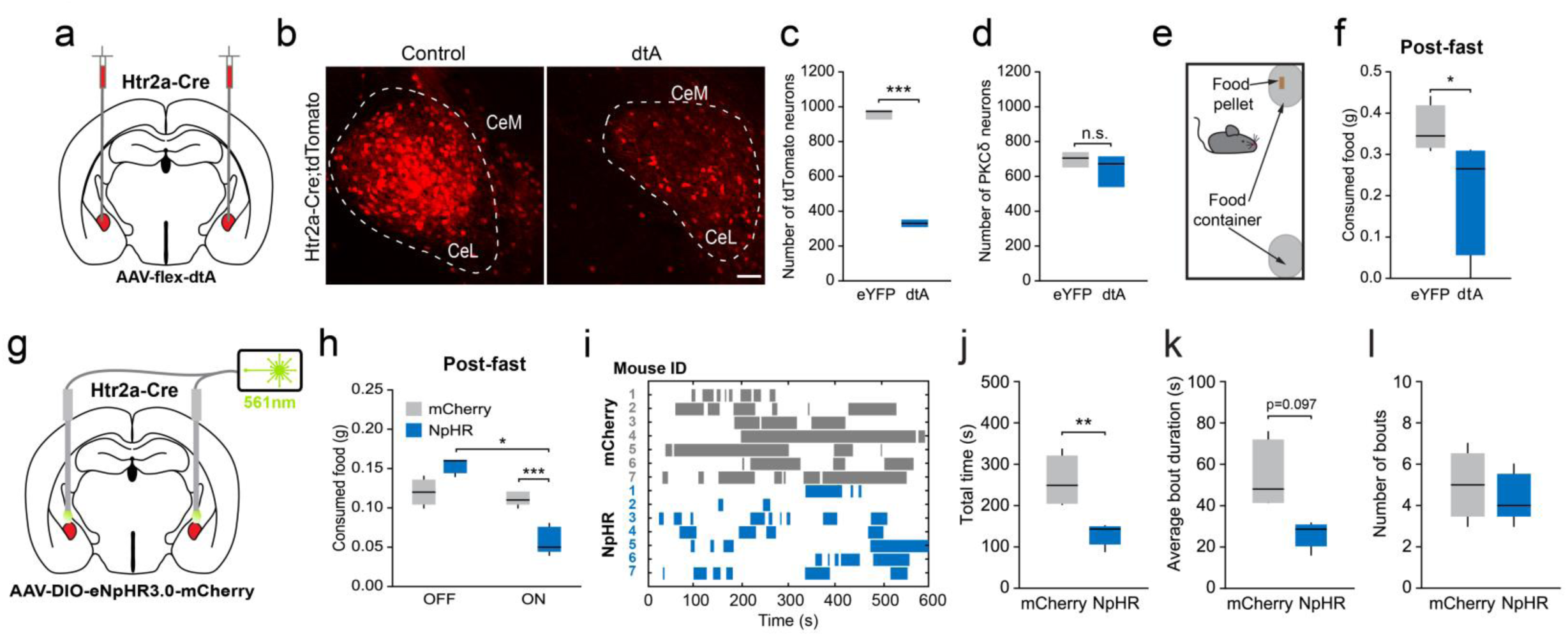
CeA^Htr2a^ neurons are required for normal food consumption. **a**, Viral delivery of AAV-flex-dtA into the CeA of *Htr2a-Cre* mice. **b**, Representative images of the CeA *of Htr2a-Cre;tdTomato* mice injected with control eYFP or with dtA virus 2 months after virus injection *(tdTomato* channel only is shown). **c**, The number of tdTomato-expressing neurons in the CeA was reduced in CeA^Htr2a^::dtA mice (n = 3 sections/ 3 mice, two-tailed unpaired t-test: t(4) = 16.88, p = <0.0001). **d**, The number of immunoreactive CeA^PKCδ^ neurons was not affected in CeA^Htr2a^::dtA mice (n = 3 sections/ 3 *mice*, two-tailed unpaired t-test: t(4) = 0.72, p = 0.5121). **e**, Scheme of free-feeding assay. **f**, CeA^Htr2a^::dtA cell ablated mice consumed less food after a 24 hour fast (n = 6 per group, two-tailed unpaired t-test: t(10) = 2.37, p = 0.0394). **g**, Optic fiber placement above CeA^Htr2a^ ::eNpHR -expressing neurons. **h**, Quantity of food consumed by food-deprived CeA^Htr2a^::eNpHR and control mice during photoinhibition and non-photoinhibited 10 minute epochs (n= 7 per group, mCherry vs NpHR ON, two-tailed unpaired t-test: t(12) = 5.20, p = 0.0002; NpHR OFF vs ON, two-tailed paired t-test: t(6) = 3.12, p = 0.0207). **i**, Raster plot of feeding bouts of example individual mice. **j**, Photoinhibition of CeA^Htr2a^ neurons decreased the total time food-deprived mice spent feeding compared to controls (n = 7 per group, two-tailed unpaired t-test: t(12) = 3.08, p = 0.0095). **k,l**, The average duration of the feeding bouts (**k;** n = 7 per group, t(12) = 1.80, p = 0.0971) and the number of bouts (**l;** n = 7 per group, t(12) = 0.24, p = 0.8142) were not significantly decreased. Box–whisker plots display median, interquartile range and 5th–95th percentiles of the distribution. *p<0.05, ***p<0.001. Scale bar: 100µm.

We next asked whether activation of CeA^Htr2a^ neurons could increase consumption under anorexigenic conditions where motivation to find and consume food is low ^21–24^. Indeed, chemogenetic activation of CeA^Htr2a^ neurons in fasted mice decreased the appetite suppressant effects of lithium chloride (LiCl) and lipopolysaccharide (LPS) that mimic toxic foods and bacterial infections respectively^25^ (Fig. 1i). Activation of CeA^Htr2a^ neurons also rescued the effect of quinine-spiked bitter food which normally reduces food intake^26^, without affecting the sensitivity of the mice to bitter tastants (Supplementary Fig. 2l-m). Together, these data demonstrate that CeA^Htr2a^ neurons promote food consumption even in the absence of physiological need and under conditions when the motivation to consume food is low.

We further examined whether activation of CeA^Htr2a^ neurons would also increase the effort made to obtain food by assessing the behavior of CeA^Htr2a^::hM3Dq mice in a food-seeking progressive-ratio task. Here, active nose-pokes were rewarded with a food pellet on a progressive-ratio 2 (PR2) schedule. We compared the performance of CeA^Htr2a^::hM3Dq mice in two consective sessions where either CNO or saline was injected prior to the experiment. CNO-treated CeA^Htr2a^::hM3Dq did not show a difference in the number of active nose-pokes or the number of consecutive nose-pokes made to obtain a single pellet (breakpoint) compared to their performance after saline treatment (Fig. 2 j and Supplementary Fig. 2o). Thus, activation of CeA^Htr2a^ neurons evokes increased consumption without affecting the motivation to work for food.

We additionally found that anxiety-like and locomotor behaviors of CeA^Htr2a^::hM3Dq mice were not significantly different from controls (Supplementary Fig. 2p-s), suggesting the altered consummatory behavior is unlikely to result from altered locomotion.

We confirmed our findings using an optogenetic approach where we expressed Cre-dependent channelrhodopsin (ChR2-eYFP) or eYFP selectively in CeA^Htr2a^ neurons and implanted optical fibers bilaterally above the CeA for somata photostimulation (Fig. 1 k and Supplementary Fig. 3a-d). Photostimulation of these neurons at 20 Hz increased food intake in satiated mice, mainly by extending the duration of eating bouts (Fig. 1l-p and Supplementary Fig. 3e-g). During the photostimulation epoch, we observed the CeA^Htr2a^::ChR2 mice engaging in stereotyped appetitive feeding-related motor behaviors, including licking of the walls and floor and chewing of the food container. Often, the mice made an analogous movement to that of holding and chewing food independent of the food pellet (Supplementary Fig. 3h and Supplementary Video 1).

The above findings demonstrate that activation of CeA^Htr2a^ neurons promoted consummatory behavior directed at food in the absence of homeostatic deficit. To explore whether CeA^Htr2a^ neurons are necessary for long-term control of food intake and body weight, we specifically ablated these neurons using a diphtheria toxin-expressing AAV (dtA) ^27^ (Fig. 2a). This resulted in loss of the majority of CeA^Htr2a^ neurons, whereas the number of CeA^PKCδ^ neurons remained unchanged (Fig. 2a-d and Supplementary Fig. 4a). Ablation of CeA^Htr2a^ neurons did not significantly affect daily food intake or body weight when the mice were maintained on a chow diet, revealing that these neurons likely do not play a role in long-term energy homeostasis, although incomplete ablation may also account for this finding (Supplementary Fig. 4b, c). However, when mice were deprived of food for 24 hours, CeA^Htr2a^ ablated mice consumed significantly less food in a free-feeding assay even though control mice were highly motivated to eat. (Fig. 2e, f). This suggests that these neurons are necessary when motivation to consume food is high. Anxiety-like and locomotor behaviors of CeA^Htr2a^::dtA mice were unaffected (Supplementary Fig. 4d-g), further supporting our findings from chemogenetic activation experiments, and suggesting the reduction in feeding is unlikely to be due to secondary effects on locomotion. To corroborate these findings, we acutely inhibited the neurons using Cre-dependent halorhodopsin (eNpHR3.0-mCherry) (Fig. 2g and Supplementary Fig. 4h, i). During the photoinhibition epoch, hungry CeA^Htr2a^::eNpHR mice consumed significantly less food than controls and food consumption was reduced compared to consumption in the absence of photoinhibition (Fig. 1h-l and Supplementary Fig. 4 j). Together, these results illustrate a role for CeA^Htr2a^ neurons in modulating food consumption.

### CeA^Htr2a^ neuron activity increases sustainably throughout eating

To further understand how CeA^Htr2a^ neurons modulate food intake, we performed *in vivo* Ca^2+^ imaging. It was previously shown that hypothalamic hunger-promoting AgRP neurons rapidly decrease their activity upon the sensory detection and consumption of food ^28–30^, whereas GABAergic neurons in the lateral hypothalamus increase activity during food seeking and consumption ^31^. To investigate the neural dynamics of CeA^Htr2a^ neurons during feeding, we performed *in* vivo Ca^2+^ imaging at single cell resolution in freely behaving mice. Here, Cre-dependent GCaMP6s was virally expressed in CeA^Htr2a^ neurons, a GRIN lens implanted above the CeA, and neuronal activity monitored when food was freely accessible using a head-mounted miniaturized microscope ^32,33^ (Fig. 3a-d and Supplementary Fig. 5a). When examining CeA^Htr2a^ neural activity during the first feeding bout after an overnight fast when food was most salient, we found that Ca^2+^ activity rapidly increased upon the start of eating and when the animals contacted the food pellet prior to consumption (Fig. 3e-g and Supplementary Fig. 5b, c and Supplementary Video 2). We then examined all feeding bouts during the imaging session and classified each neuron by comparing the average activity during each bout to the proceeding inter-bout interval (Supplementary Fig. 5d). This revealed that a subset of the CeA^Htr2a^ neurons consistently increased activity during eating (22%) (Fig. 3h-k and Supplementary Fig. 5e) while there were some that reduced their activity (10%) (Fig. 3 k and Supplementary Fig. 5f-h).

**Figure 3.**
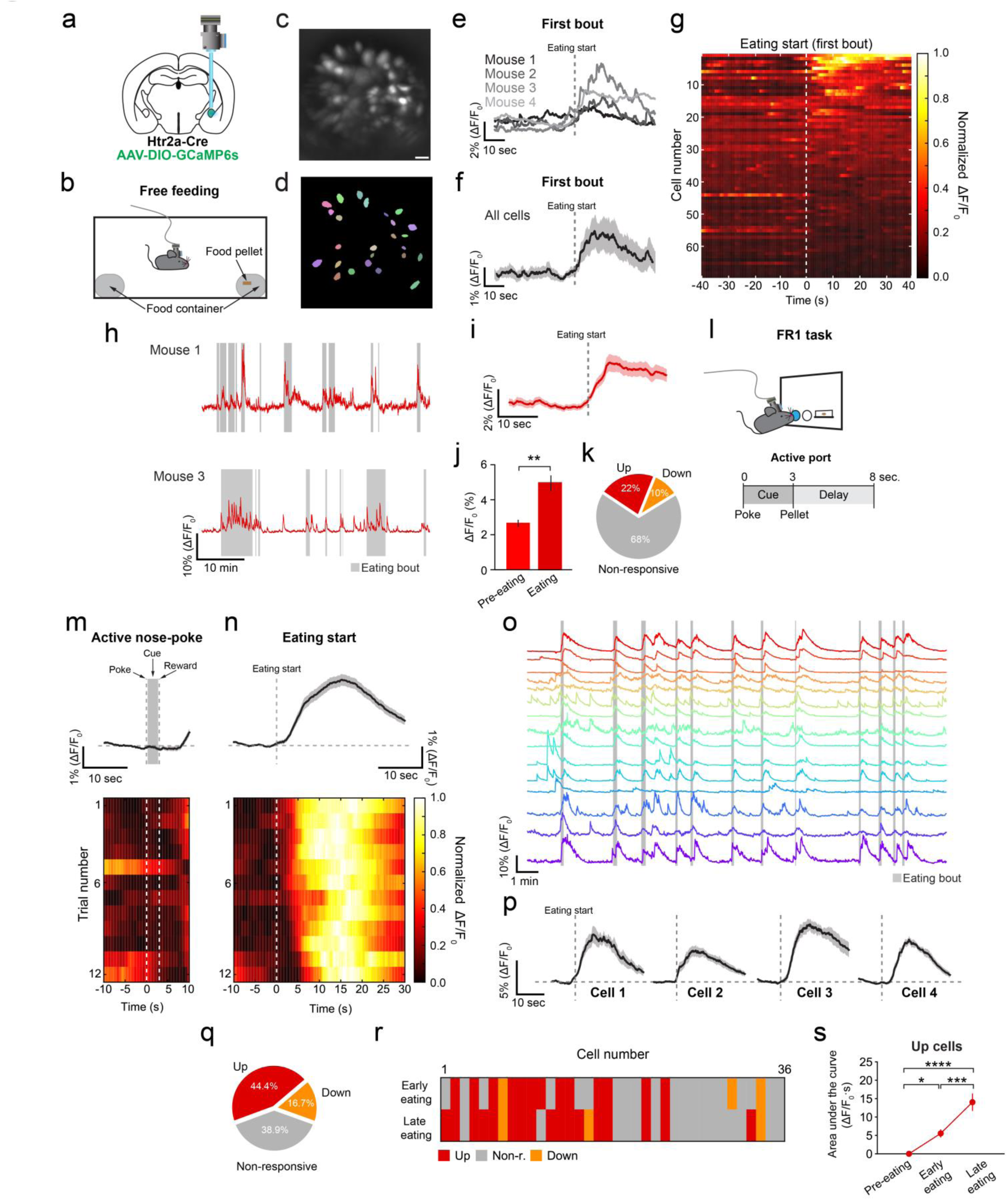
CeA ^Htr2a^ neurons increase activity during food consumption. **a**, Position of the GRIN lens above GCaMP6s-expressing CeA*^Htr2a-Cre^* neurons. **b**, Scheme of free-feeding assay. **c,d**, Maximum projection image of a representative imaging plane of CeA^Htr2a^::GCaMP6s neurons (**c**) and corresponding region of interest (ROI) cell masks (**d**). **e**, Mean Ca^2+^ responses of cells from individual animals aligned to the first eating bout onset after an overnight fast (n = 4 mice). **f**, Mean Ca^2+^ response of all recorded cells (n = 69 across 4 mice) aligned to food eating onset of the first bout. Shading = SEM. **g**, Heat map of normalized Ca^2+^ responses from individual CeA ^Htr2a^ ::GCaMP6s neurons prior to and during the first eating bout. Vertical stippled white line = eating onset. **h**, Example Up cells from two mice. Grey bars = eating bouts. **i, j**, Mean Ca^2+^ responses of cells classified as Up (n = 15/69) before and upon eating onset of all bouts in a 40 minute period (n = 15 cells across 4 mice, two-tailed paired t-test: t(14) = 3.12, p = 0.0023). **k**, Classification of 69 cells based on preference index. **l**, Scheme of FR1 task. **m,n**, Population Ca^2+^ responses of all recorded neurons (n = 36 cells across 3 mice, 12 trials) aligned to active nose-poke (**m**) and eating start (**n**). Upper = mean. Lower = heat map of normalized Ca^2+^ responses from all neurons. Each row is one trial. **o**, Ca^2+^ responses of individual neurons from a representative mouse during the FR1 task. Grey bars = eating bouts. **p**, Ca^2+^ responses of representative cells averaged across 12 trials. **q**, Classification of 36 cells during the eating bout. **r**, Heat map of cell classifications during each half of the eating bout. Activity of 36% of neurons increased during the early phase of eating, compared to 47% during the late phase. **s**, Mean activity of Up cells before eating onset and during each half of the eating bout (n = 16 cells across 3 mice, one-way repeated measures ANOVA: Time-point: F(2,15) = 28.53, p < 0.0001, Bonferroni post-hoc analysis: *p<0.05, ***p<0.001, ****p<0.0001). Bar graphs indicate mean ± SEM. *p<0.05. **p<0.01. ***p<0.001. ***p<0.0001. Scale bar: 20µm.

Next, we examined the dynamics of CeA^Htr2a^ neurons in more detail to determine whether CeA^Htr2a^ neurons encoded the appetitive and/or consummatory aspects of food intake. To do so, we recorded the activity of CeA^Htr2a^::GCaMP6s neurons during a fixed ratio-1 (FR1) task, where food restricted mice were trained to nose-poke for a 20mg food pellet reward (Fig. 3l), such that the appetitive and consummatory phases of food intake were well defined. The results revealed that CeA^Htr2a^ neurons did not increase their activity during the appetitive phase, i.e. the active nose-poke and cue phases that signaled reward delivery (Fig. 3 m), but during food consumption with activity increasing upon food contact (Fig. 3 n, Supplementary Fig. i) (Supplementary Video 3). Upon examining the response profiles of individual cells, we found that the activity profiles were consistent throughout the 12 recorded FR1 trials, but that the onset of the cells varied (Fig. 3o,p), with some cells exhibiting a rise in Ca^2+^ activity just prior to or time-locked to eating start, while other cells increased activity with a delayed onset (Fig. 3p, Supplementary Fig. 5 j). Like the free-feeding task, we also found that a subset of neurons were active during eating (44% of all cells) (Fig. 3q). Classification of the cells during the first and the second half of the eating bout revealed that some initially non-responder cells exhibited a late-onset increase in activity in the second half of the eating bout (36% active cells in the first half versus 47% in the second half) (Fig. 3r). This suggests that during eating, CeA^Htr2a^ neurons were recruited to the active ensemble and that overall the activity of CeA^Htr2a^ neurons increased during eating (Fig. 3s and Supplementary Fig. 5k). Together these data revealed that during the process of food seeking and eating in different behavior contexts, CeA^Htr2a^ neurons are consistently and specifically active during food consumption, suggesting that ongoing activity of these neurons may propagate eating behavior.

### Activity of CeA^Htr2a^ neurons is positively reinforcing

Given that food is an innately rewarding stimulus and that CeA^Htr2a^ neurons are active during its consumption, these neurons may potentiate eating by positively reinforcing eating behavior. If this hypothesis was correct, mice should seek out activation of CeA^Htr2a^ neurons. We therefore assessed the valence of optogenetic activation of CeA^Htr2a^ neurons. In a real-time place preference assay (RTPP)^34^ (Fig. 4a), CeA^Htr2a^::ChR2 mice exhibited significant place preference for the photostimulation-paired chamber, relative to controls (Fig. 4b, c; Supplementary Fig. 6a, b) while locomotor behavior in the photostimulation-paired chamber was not affected (Supplementary Fig. 6c, d). To assess if mice would perform instrumental responses for CeA^Htr2a^ neuron activation, we trained mice to nose-poke for intracranial optical self-stimulation with a fixed-ratio one schedule of reinforcement (Fig. 4d). CeA^Htr2a^::ChR2 mice nose-poked to receive 20Hz photostimulation significantly more than controls (Fig. 4e, f and Supplementary Video 4). These results establish that CeA^Htr2a^ neuron activity is intrinsically positively reinforcing. As food consumption is, in part, influenced by its rewarding properties such as taste and palatability, we next assessed whether this positive valence signal increases food consumption by enhancing the rewarding properties of food. We thus investigated whether activity of CeA^Htr2a^ neurons could condition preference to specific flavors (Fig. 4g). Here, CeA^Htr2a^::ChR2 mice were allowed to consume two differently flavored non-nutritive gels. After baseline preference was determined, the mice were conditioned in alternating sessions, where the less-preferred flavor was paired with CeA^Htr2a^ photostimulation. After conditioning, the mice were simultaneously exposed to both flavors. We found that concurrent CeA^Htr2a^ photostimulation reversed the initial flavor preference, such that the initially less preferred gel became more preferred (Fig. 4h). Thus, the positive valence signal conveyed by CeA^Htr2a^ neurons modulates food consumption by influencing the rewarding properties of food.

**Figure 4.**
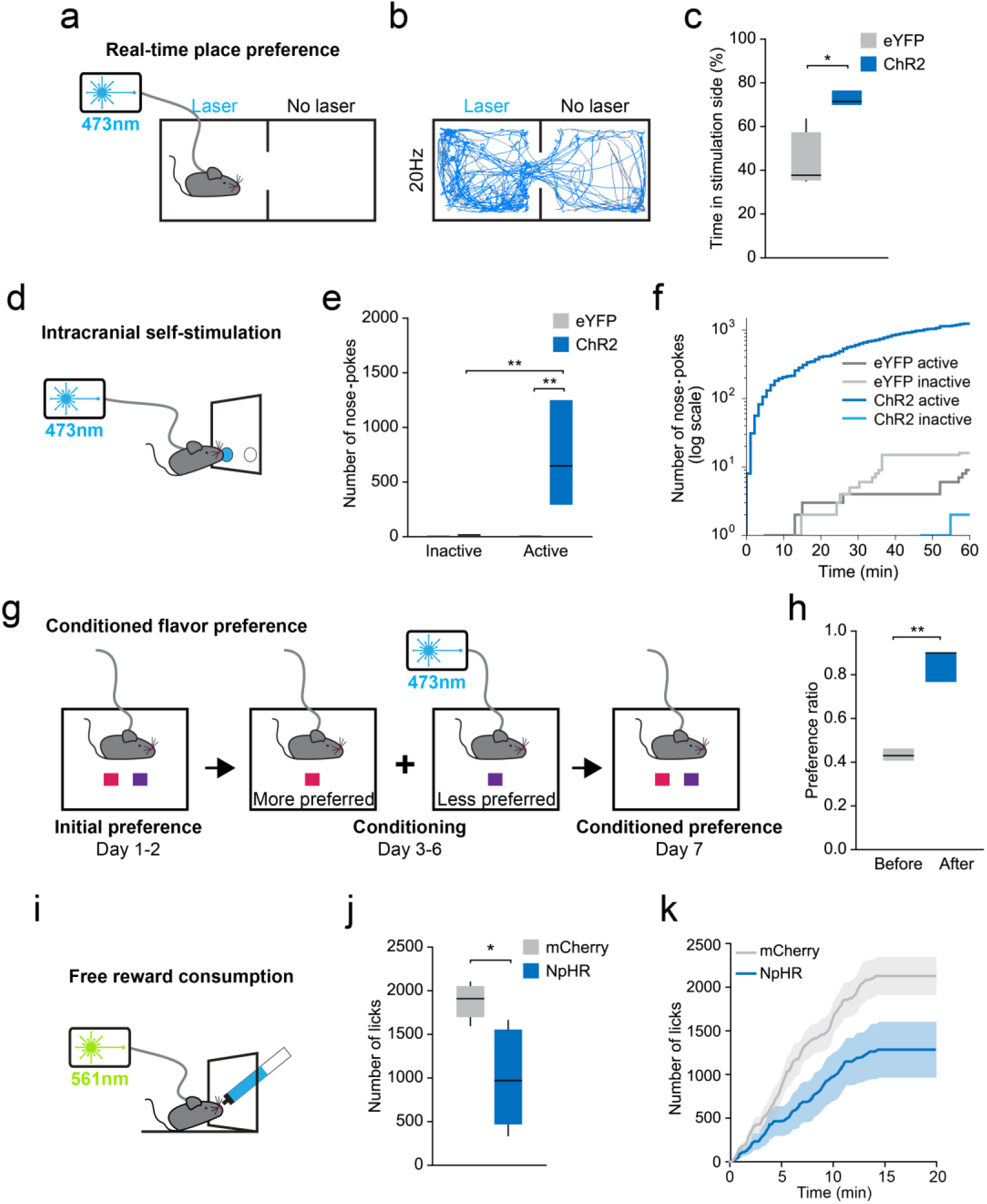
CeA^Htr2a^ neuron activity is positively reinforcing and modulates reward consumption. **a**, Scheme depicting real-time place preference (RTPP) paradigm where the side of the chamber marked ‘Laser’ was paired with 473nm 20 Hz photostimulation of CeA^Htr2a^ neurons. **b**, Representative locomotor trace of a CeA^Htr2a^::ChR2 mouse that received 20 Hz photostimulation in the ‘Laser’ compartment. **c**, CeA^Htr2a^::ChR2 mice spent more time in the photostimulated side of the RTPP chamber (n = 6 per group, two-tailed unpaired t-test: t(10) = 2.80, p = 0.0192). **d**, Scheme depicting intracranial self-stimulation (ICSS) paradigm. The active port for 20 Hz optogenetic self-stimulation of CeA^Htr2a^ neurons is indicated in blue. **e**, Number of nose-pokes of CeA^Htr2a^::ChR2 mice at the active and inactive port during the 60 minute session (n = 5 per group: two-way ANOVA: Virus: F(1,16) = 8.77, p = 0.0092, Nosepoke: F(1,16) = 8.30, p = 0.0109, Interaction: F(1,16) = 8.48, p = 0.0102, Bonferroni post-hoc analysis: **p<0.01). **f**, Cumulative nose-poke responses made by representative CeA^Htr2a^::ChR2 and CeA^Htr2a^::eYFP control mice. **g**, Scheme of conditioned flavor preference experiment where the less-preferred of two different flavoured gels was paired with photostimulation of CeA^Htr2a^ neurons. **h**, Photostimulation of CeA^Htr2a^ neurons reversed flavour preference of the initially less-preferred flavor (n = 5, two-tailed paired t-test: t(4) = 4.99, p = 0.0076). **i**, Scheme depicting free consumption of a liquid palatable reward paradigm concurrent with constant 561nm photoinhibition of CeA^Htr2a^ neurons. **j**, Photoinhibition of CeA^Htr2a^ neurons in *ad libitum* fed mice decreased licking of a spout that delivered a palatable reward (n = 7 mCherry, n = 8 NpHR, two-tailed unpaired t-test: t(13) = 2.38, p = 0.0333). **k**, Mean cumulative licks of palatable reward made by CeA^Htr2a^ ::NpHR and CeA^Htr2a^::mCherry controls over the 20 minute session. Shading = SEM. Box–whisker plots display median, interquartile range and 5th–95th percentiles of the distribution.*p<0.05, **p<0.01. Scale bars: 100µm.

We further explored this notion by asking whether CeA^Htr2a^ neurons modulate food palatability. To test this, we trained CeA^Htr2a^::NpHR mice to lick a spout for a palatable liquid reward (Fig. 4i). Mice were tested for licking behavior during constant photoinhibition in the *ad lib* fed state, where consumption is driven by palatability rather than homeostatic need. Photoinhibited CeA^Htr2a^::NpHR mice consumed less of the reward than mCherry-expressing mice during the minute photoinhibition period (Fig. 4 j,k) which was not observed in hungry animals (Supplementary Fig. 6j). Interestingly, we found that CeA^Htr2a^::NpHR mice did not avoid the photoinhibited side of the chamber in the RTPP assay (Supplementary Fig. 6e-g) and displayed similar locomotor behavior to controls (Supplementary Fig. 6h-i). Thus, silencing of CeA^Htr2a^ neurons does not lead to induction of intrisinc aversion, but leads to decreased consumption as eating is no longer positively reinforced. Together, these data show that activity of CeA^Htr2a^ neurons is reinforcing and that it modulates food reward, suggesting that these neurons function to promote ongoing eating behavior through a positive valence signal.

### Antagonism between CeA neural modulators of food consumption

Given that CeA^Htr2a^ neurons promote food consumption and reside alongside CeA^PKCδ^ neurons that suppress feeding, we investigated the local circuit interaction of these neuronal populations to understand how the CeA bidirectionally modulates food consumption. To determine whether CeA^Htr2a^ neurons could inhibit CeA^PKCδ^ neurons, we targeted CeA^Htr2a^ neurons with Cre-dependent AAV-ChR2 in *Htr2a-Cre;tdTomato* mice (Fig. 5a, b). Whole-cell recordings of CeA tdTomato-negative neurons revealed light-evoked, short latency, picrotoxin-sensitive inhibitory postsynaptic currents (IPSCs) in all recorded neurons (Fig. 5c, d). Post-hoc identification of neurobiotin filled recorded neurons revealed that 50% were PKCδ+ (Fig. 5e). Paired recordings in acute slices from *Htr2a-Cre;tdTomato* mice revealed unidirectional connections between cell pairs with reversal potentials typical for GABA receptors. (Supplementary Fig. 7a-c). Together, these results suggest strong monosynaptic inhibition from CeA^Htr2a+^ onto CeA^Htr2a^‒ neurons including CeA^PKCδ^ neurons.

**Figure 5.**
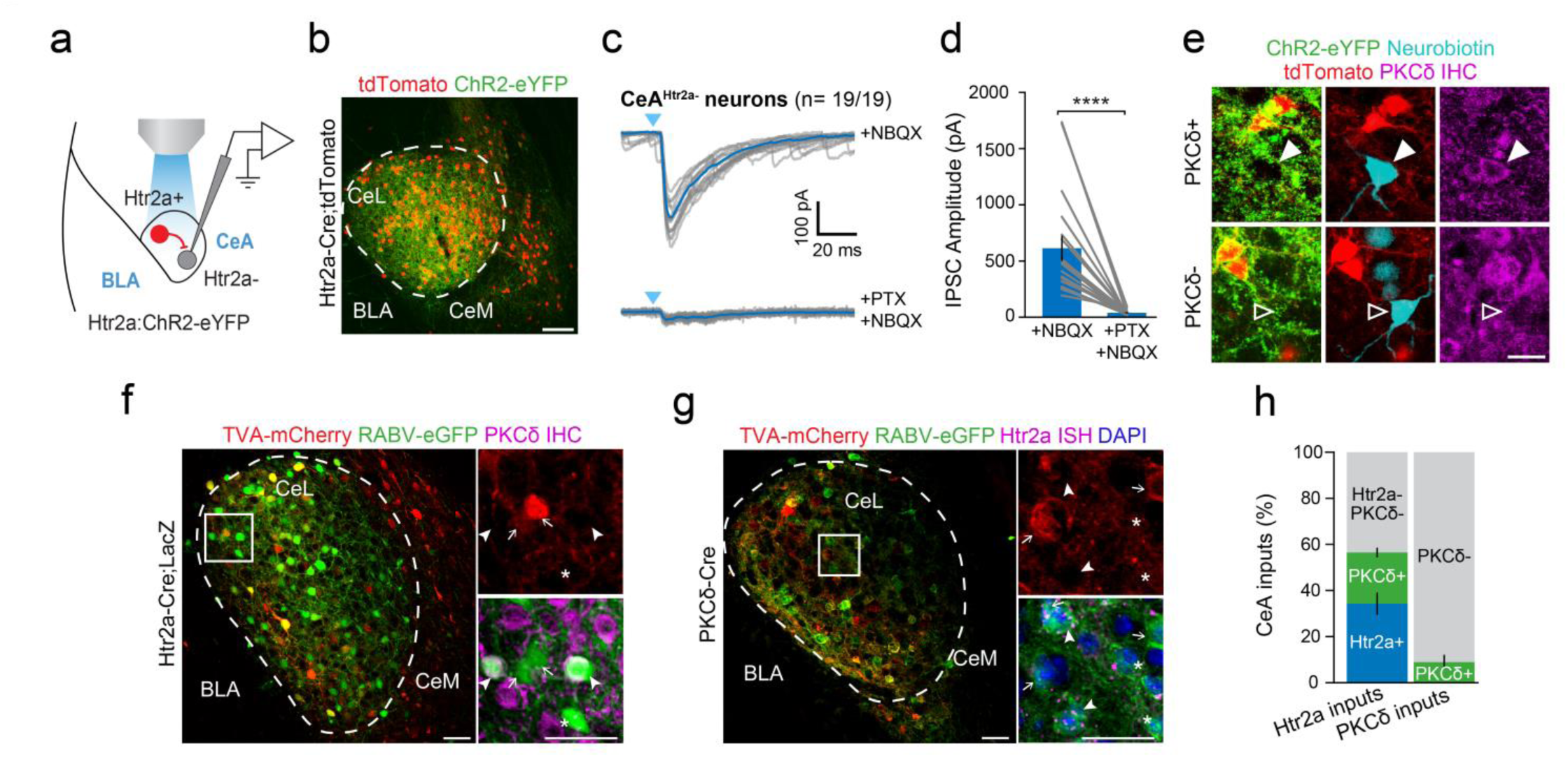
CeA neuronal modulators of feeding behavior form reciprocal inhibitory connections. **a**, Scheme of whole-cell recording from *CeA^Htr2a-^* neurons in *Htr2a-Cre;tdTomato* slices transduced with ChR2-eYFP. **b**, Representative image of ChR2-eYFP and tdTomato expression in the CeA of a *Htr2a-cre;tdTomato* mouse. **c**, Overlay of light evoked (blue triangle, 1ms) individual (grey) and the average (blue) IPSCs in the absence (top) and presence of PTX (bottom) in postsynaptic *CeA^Htr2a^-^Cre;tdTomat^o-* neurons (average onset latency of 2.7 ±0.05 ms). All recordings include NBQX to block excitatory synaptic transmission. **d**, Quantification of light evoked responses (mean IPSC amplitude 604.93 ±106.04 pA) (n = 19 Htr2a+ cells, two-tailed paired t-test: t(18) = 5.42, p < 0.0001). **e**, Examples of recorded and neurobiotin-filled CeA*^Htr2a-Cre;tdTomato-^*neurons *post-hoc* identified as PKCδ+ (3/6 neurons) and PKCδ-. Filled and empty arrowheads show PKCδ+ and PKCδ- neurons, respectively. **f,g**, Identification of local monosynaptic inputs to CeA^Htr2a^ (**f**) or CeA^PKCδ^ neurons **(g)** using Cre-dependent, rabies virus-based monosynaptic tracing. White boxes indicate the location of the high-magnifications on the right. Arrows indicate starter cells (co-labeled with eGFP and mCherry). Arrowheads indicate input neurons (RABV-eGFP+ cells) immunolabeled for PKCδ antibody **(f)** or *in situ* hybridized for Htr2a mRNA probe **(g)**. Asterisks denote RABV-eGFP-labeled input neurons only. **h**, Quantification of the relative abundance of PKCδ+ and Htr2a+ input cells to CeA^Htr2a^ and CeA^PKCδ^ neurons (n= 3 *Htr2a-Cre* and *PKCδ-Cre* mice). Bar graph values are mean ± SEM. Scale bars: 50µm.

To anatomically map local synaptic inputs to CeA^Htr2a^ and CeA^PKCδ^ neurons within the CeA, we performed Cre-dependent, rabies virus-based monosynaptic retrograde tracing^35^ paying attention that rabies-mediated labelling of input neurons was dependent on Cre expression (Supplementary Fig. 7d). We injected *Htr2a-Cre;LacZ* or *PKCδ-Cre* mice with Cre-dependent AAVs that express the avian EnvA receptor (TVA) and rabies virus envelope glycoprotein (RG) in combination with a modified rabies virus SAD∆G-EGFP (EnvA) (Supplementary Fig. 7e, f). These experiments revealed that local inputs to CeA^Htr2a^ neurons came from CeA^PKCδ^ neurons and CeA^Htr2a^ neurons in similar proportions (Fig. 5f, h and Supplementary Fig. 7g). In addition, we confirmed that CeA^PKCδ^ neurons received monosynaptic inputs from CeA^Htr2a^ neurons (Fig. 5g) and showed that only a small portion of monosynaptic inputs to CeA^PKCδ^ neurons originated from CeA^PKCδ^ cells (Fig. 5h and Supplementary Fig. 7h). In summary, these results revealed that CeA^Htr2a^ and CeA^PKCδ^ neurons form monosynaptic reciprocal connections within the CeA.

### Inhibition of PBN by CeA^Htr2a^ neurons is positively reinforcing and modulates food consumption

To elucidate the neurocircuitry by which CeA^Htr2a^ neurons modulate food consumption, we anatomically mapped the long-range innervation fields of these neurons. By expressing a Cre-dependent, synaptically targeted fluorophore (AAV-Synaptophysin-myc) or mCherry selectively in CeA^Htr2a^ neurons (Fig. 6a), we observed dense efferent fields most prominently in the PBN (Fig. 6b, Supplementary Fig. 8a, b and Supplementary Video 5). Given the described role of the PBN in appetite suppression^11,17,36–38^ we hypothesized that CeA^Htr2a^ neurons promote feeding by suppressing PBN neurons. To further investigate the cell type specificity of this projection, we injected retrogradely transported beads into the PBN of *Htr2a-Cre;tdTomato* mice (Fig. 6c) and observed that 67% of retrobead-positive neurons were CeA^Htr2a^ neurons, while less than 1% were PKCδ immunopositive (Fig. 6d, e). To assess the functionality of this projection we injected Cre-dependent ChR2-eYFP into the CeA of *Htr2a-Cre* and *PKCδ-Cre* mice (Fig. 6f) and performed whole-cell recordings from PBN neurons inside the area of ChR2 innervation (Fig. 6f, g). In slices from *Htr2a-Cre* animals, we detected light evoked, short latency, picrotoxin-sensitive IPSCs in most recorded PBN neurons, whereas in *PKCδ-Cre* animals very few cells responded and only with very small amplitudes (<30 pA) (Fig. 6h, i). Light stimulation also suppressed evoked firing by current-injection in PBN neurons in a picrotoxin-sensitive manner (Fig. 6j and Supplementary Fig. 8c). Post-hoc identification of neurobiotin filled neurons revealed that 12 out of 14 neurons were negative for CGRP, a marker of anorexia-promoting neurons^17^ (Supplementary Fig. 8d). Although the molecular identity of PBN neurons engaged by CeA^Htr2a^ neurons remains to be elucidated, the strength of the CeA^Htr2a^-PBN connection implicates this projection in regulating the function of the PBN.

**Figure 6.**
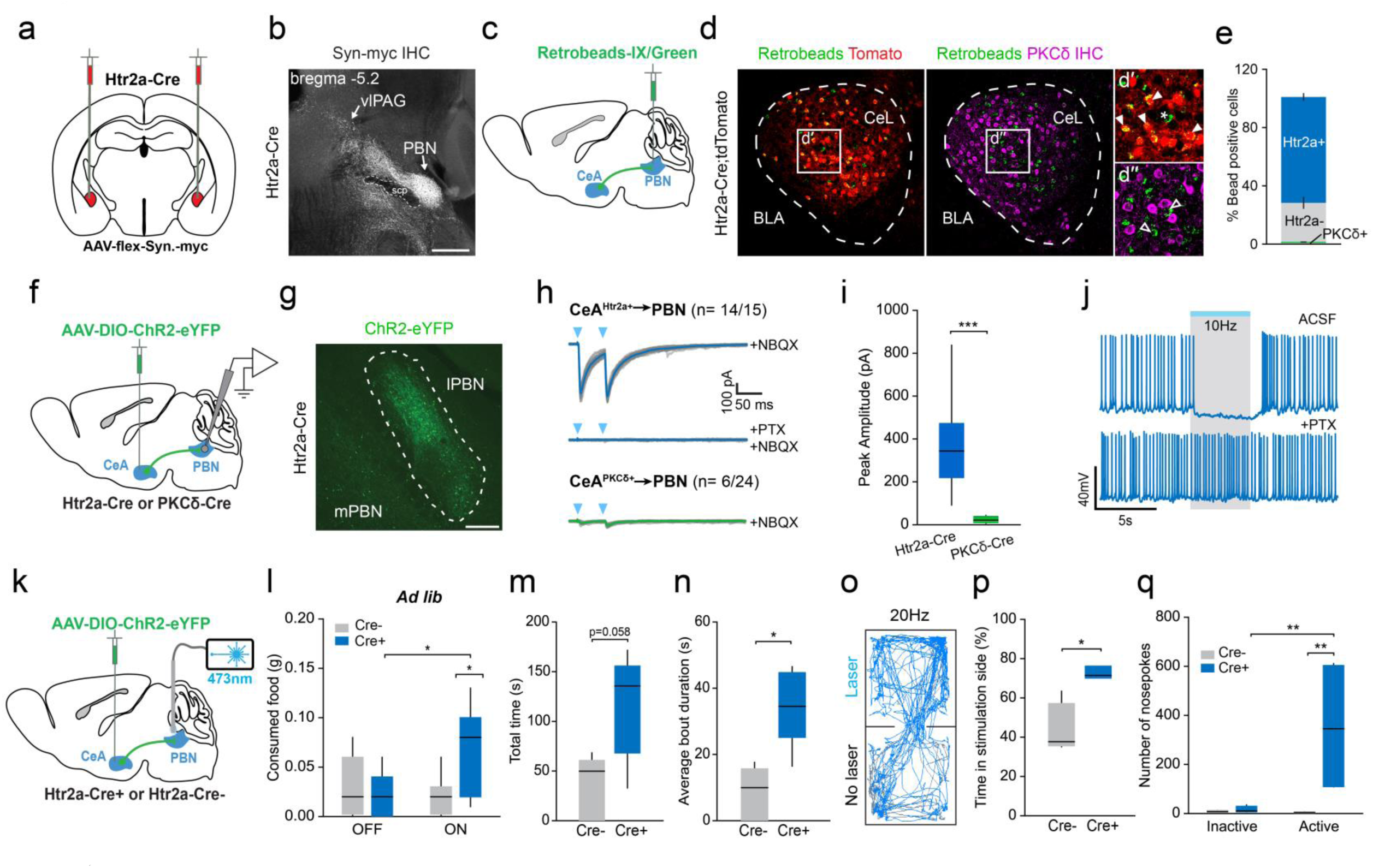
CeA^Htr2a^ neuron inhibition of the PBN is rewarding and modulates consummatory behaviour. **a**, Viral delivery of Synaptophysin-myc into the CeA of *Htr2a-Cre* mice. **b**, Anti-myc immunohistochemistry reveals CeA^Htr2a^ axon terminals densely innervating the PBN. **c**, Scheme of retrograde-tracing strategy from the PBN. **d, e**, Retrobeads retrogradely transported from the PBN predominately colocalize with *Htr2a-Cre+* neurons in the CeA. Closed arrowheads= bead+/tdTomato+, asterisk= bead+/ tdTomato- in **d’**, open arrowheads= bead+ / PKCδ- in **d’’** (n=3 mice). **f**, Viral targeting of CeA to investigate the CeA→PBN projection. **g**, ChR2-expressing axons of CeA^Htr2a^ neurons terminate in the lateral PBN. **h, i**, Photoactivation of CeA^Htr2a^ terminals in the PBN induce high amplitude PTX-sensitive IPSCs in 14/15 PBN neurons, while activation of CeA^PKCδ^ terminals induced weak responses in 6/24 recorded PBN neurons (Mann Whitney test: p = 0.0008). All recordings include NBQX to block excitatory synaptic transmission. **j**, Photostimulation of CeA^Htr2a^ terminals by 473nm light pulses suppressed induced firing of PBN neurons by current injection in a PTX-dependent manner. **k**, Virus injection of ChR2-eYFP bilaterally into the CeA with bilateral fiber optic implant over PBN (only one side is shown). **l**, Quantity of food consumed by *ad libitum* fed CeA^Htr2a^::ChR2→PBN and control mice during 20 Hz photoactivation and non-photoactivated 20 minute epochs (n = 10 Cre-, n = 9 Cre+ Cre- vs Cre+ ON, two-tailed unpaired t-test: t(17) = 2.39, p = 0.0289; Cre+ OFF vs ON, two-tailed paired t-test: t(8) = 2.38, p = 0.0449). **m**, Photostimulation of CeA^Htr2a^::ChR2 terminals in the PBN at 20 Hz did not significantly increase the total time spent eating (n = 7 per group, two-tailed paired t-test: t(12) = 2.09, p = 0.0584). **n**, The average duration of eating bouts was increased in CeA^Htr2a^::ChR2 mice (n = 7 per group, Mann Whitney test: p = 0.0330). **o**, Example locomotor trace of a mouse in the RTPP experiment where the ‘Laser’ side was paired with 20 Hz photostimulation of CeA^Htr2a^ presynaptic terminals in the PBN. **p**, Time spent by CeA^Htr2a^::ChR2 → PBN and control mice in the photostimulated side of the RTPP chamber (n = 6 Cre-, n = 5 Cre+, two-tailed unpaired t-test: t(9) = 2.50, p = 0.0338). **q**, Number of nose-pokes at the active and inactive ports made by CeA^Htr2a^::ChR2 → PBN mice and control mice during an ICSS session (n = 6 per group, two-way ANOVA: Virus: F(1,20) = 6.40, p = 0.0199, Nosepoke: F(1,20) = 6.11, p = 0.0225, Interaction: F(1,20) = 5.91, p = 0.0246, Bonferroni post-hoc analysis: **p<0.01). Box-whisker plots display median, interquartile range and 5th-95th percentiles of the distribution.Bar graphs indicate mean ± SEM. *p<0.05, **p<0.01, ***p<0.001. Scale bars: 100µm.

Hence, we sought to determine whether inhibition of this region by CeA^Htr2a^ neurons was sufficient to promote CeA^Htr2a^ neuron-mediated feeding-related behaviors. We transduced CeA^Htr2a^ neurons with Cre-dependent ChR2-eYFP and placed optic fibers bilaterally above the PBN (Fig. 6 k and Supplementary Fig. 8e,f). We found that photostimulation of the CeA^Htr2a^ presynaptic terminals in the PBN elicited a modest yet significant increase in food intake in *ad libitum* fed CeA^Htr2a^::ChR2 mice. (Fig. 6l-n and Supplementary Fig. 8g-i). We also probed whether activation of the CeA^Htr2a^ → PBN projection was intrinsically rewarding. Indeed, 20Hz photostimulation of the CeA^Htr2a^ presynaptic terminals in the PBN elicited significant place preference (Fig. 6o-p) and increased nose-poking behavior for optogenetic self-stimulation (Fig. 6q). Together, our results suggest that the elicitation of food consumption and positive reinforcement properties of CeA^Htr2a^ neurons are in part mediated through inhibition of neurons in the PBN.

Given that CeA^Htr2a^ neurons function in part through inhibition of neurons in the PBN and that CeA^PKCδ^ neurons were proposed to suppress feeding via local inhibition of CeA^PKCδ‒^ neurons^14^, we next determined whether CeA^PKCδ^ neurons provide inhibition onto PBN-projecting neurons. To reveal local monosynaptic inputs of PBN-projecting CeA cells, we used the TRIO strategy^39^, which combines retrograde transport of virally expressed Cre recombinase from the PBN to the CeA and Cre-dependent monosynaptic rabies tracing in the CeA (Supplementary Fig. 9a,b). This experiment revealed that a proportion of inputs to PBN-projecting CeA neurons came from CeA^PKCδ^ neurons (Supplementary Fig. 9c,d). To determine if CeA^PKCδ^ neurons inhibit CeA PBN-projecting neurons, we expressed Cre-dependent ChR2-eYFP in *PKCδ-Cre* mice and identified PBN-projecting neurons by injecting retrobeads into the PBN (Supplementary Fig. 9e). In acute slices, whole-cell recordings of bead-positive CeA neurons revealed light evoked picrotoxin sensitive IPSCs (Supplementary Fig. 9f). These results reveal a circuit by which the CeA can modulate food consumption via interactions between genetically-defined cell types.

### Inputs to CeA^Htr2a^ neurons arise from feeding relevant brain regions

The newly identified role of CeA^Htr2a^ neurons in food consumption prompted us to explore the circuits in which these neurons integrate. We extended our monosynaptic tracing analysis to identify regions providing long-distance synaptic inputs to CeA^Htr2a^. We found that CeA^Htr2a^ neurons receive direct synaptic inputs from a wide range of brain regions, including the neural components of the gustatory and visceroceptive pathway^40^ such as the insular cortex (IC),gustatory thalamus (VPMpc) and the lateral parabrachial nucleus (lPBN) as well from other feeding centers such as the hypothalamic arcuate nucleus^41^ and the parasubthalamic nucleus (PSTN)^42^ (Fig. 7a and Supplementary Fig. 10a-c). The midbrain dorsal raphe (DR) nucleus as well as the substantia nigra pars lateralis (SNL), which contain serotonergic^43^ and dopaminergic^44^ neurons, respectively, represented major inputs to CeA^Htr2a^ neurons (Fig. 7a and Supplementary Fig. 10c). Interestingly, we observed that the numbers of neurons in the input regions varied between animals (Supplementary Fig. 10d) raising the possibility that the starter cells in each experiment represented subsets of spatially clustered CeA^Htr2a^ neurons that harbor distinct input patterns. Pairwise correlation followed by hierarchical clustering analysis revealed a strong positive correlation between inputs from the IC and VPMpc, whereas hypothalamic and midbrain nuclei formed a separate cluster (Fig. 7b and Supplementary Fig. 10e). These results suggested that CeA^Htr2a^ neurons that receive information from cortical and thalamic nuclei may be distinct from those innervated by the hypothalamic and midbrain structures.

**Figure 7.**
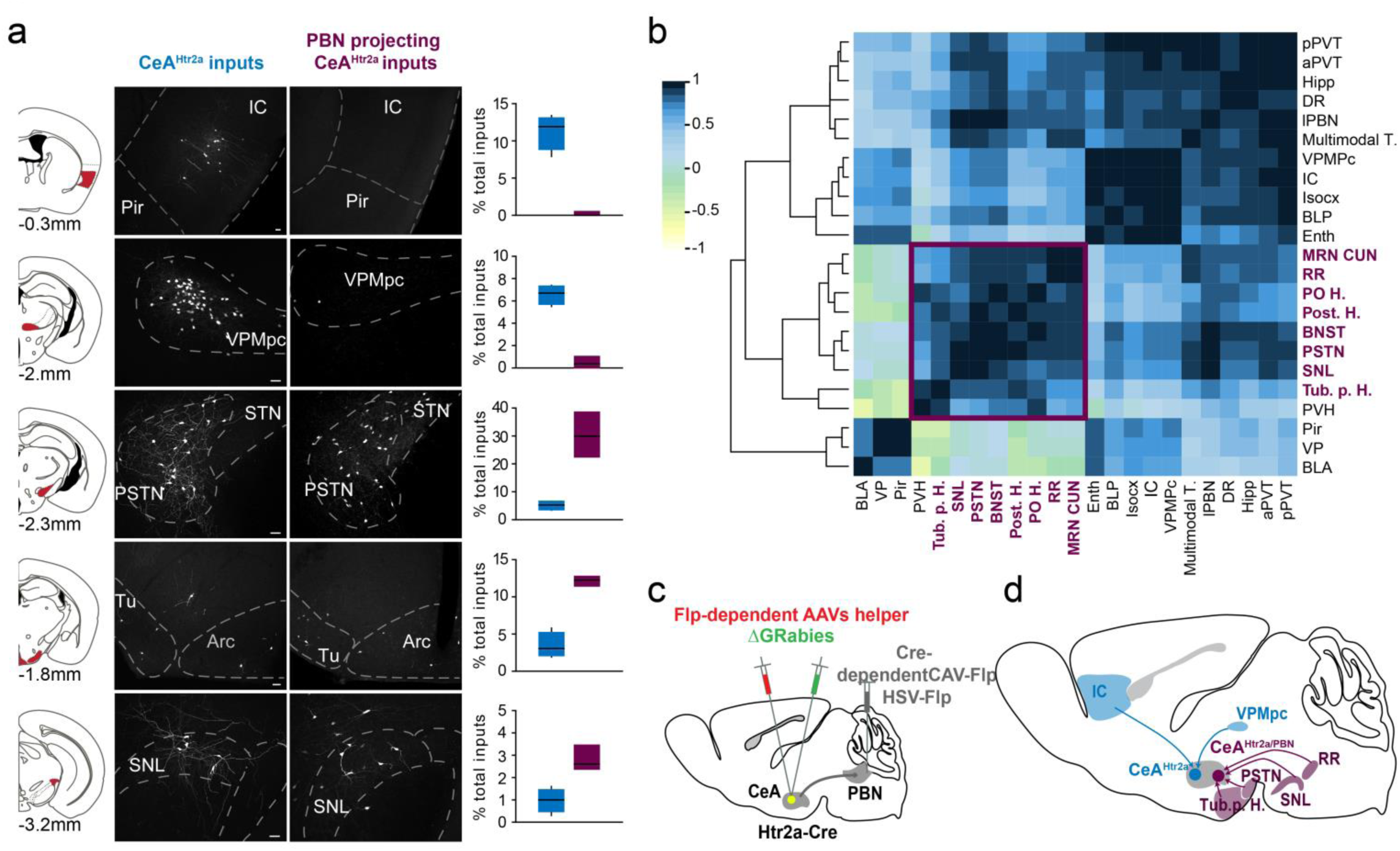
Inputs to CeA^Htr2a^ neurons come from feeding relevant brain regions. **a**, Distribution of long-distance monosynaptic inputs (RABV-eGFP+ cells) to CeA^Htr2a^ and PBN-projecting CeA^Htr2a^ cells in the IC, VPMpc, PSTN, Tub. p. hypothalamus comprising tuberal nucleus (TU) and arcuate nucleus (Arc), and SNL. Approximate coronal section planes are shown on the left with distance (anterior-posterior) from bregma. Graphs show for each region, the proportion of inputs to CeA^Htr2a^ (blue) and PBN-projecting CeA^Htr2a^ neurons (magenta), normalized against the total number of input cells in each animal. **b**, Hierarchical clustering on Pearson’s pairwise correlation coefficients between all input regions counted from 6 *Htr2a-Cre* tracing experiments. The color scale indicates the degree of correlation. The cluster of input regions to PBN-projecting CeA^Htr2a^ neurons is marked in magenta color. **c**, Schematic representation of the cTRIO strategy used to reveal monosynaptic inputs to PBN-projecting CeA^Htr2a^ neurons. **d**, Schematic summary showing examples of long distance monosynaptic inputs to CeA^Htr2a^ cells. In blue, IC and VPMpc target a distinct subset of CeA^Htr2a^ neurons. In magenta, PSTN, RR, SN and Tub. p. specifically project onto PBN-projecting CeA^Htr2a^ cells. For abbreviations of each region, see Supplementary Figure 10. Bar graph values are mean ± SEM. Scale bars: 50µm.

To investigate whether PBN-projecting CeA^Htr2a^ neurons receive a distinct subset of these inputs, we used a modified version of the TRIO strategy^39^ which utilizes retrograde transport from the PBN to the CeA of virally expressed Flp recombinase whose expression is Cre-dependent, in combination with monosynaptic rabies tracing in the CeA, under control of Flp recombinase (Fig.7c and Supplementary Fig. 10a, b). This experiment revealed that PBN-projecting CeA^Htr2a^ cells receive long range inputs from discrete brain regions including those that process information relevant for energy homeostasis (Arc)^41^, those containing neurons that respond to ingestion of palatable food (PSTN)^42^ and those that harbor dopaminergic neurons (SNL and RR)^44^.

We found these regions to be identical to the ones targeting the broader PBN-projecting CeA population (Supplementary Fig. 10c), and, in our pairwise correlation analysis, to belong to one distinct cluster (Fig. 7j). The majority of these regions represented major inputs to CeA^Htr2a^ neurons and minor inputs to CeA^PKCδ^ neurons (Supplementary Fig. 10c). In summary, we show that the CeA^Htr2a^ population receives diverse inputs from multiple brain areas known to process sensory and homeostatic information and that CeA^Htr2a^ neurons are comprised of subclusters defined by their input patterns. Additionally, those CeA^Htr2a^ neurons projecting to the PBN receive a distinct set of inputs from feeding relevant brain regions.

## Discussion

Seeking and consumption of food is goverened by a complex framework of behaviors. Our study defines the neural mechanisms by which molecularly-defined neurons within the CeA promote food consumption. Here we show that Htr2a-expressing CeA neurons increase activity during eating and that their activity bi-directionally modulates food intake. Further, we demonstrate that activation of these neurons is positively reinforcing. Together, these data support a model where CeA^Htr2a^ neuron activity leads to sustained eating behavior by both local and long-range circuit mechanisms.

Consumption of food consists of serialized motivated behaviors that constitute appetitive and consummatory actions. Our findings demonstrate that the circuits associated with CeA^Htr2a^ neurons modulate the consummatory phase of eating. Artificial activation of CeA^Htr2a^ neurons specifically increased food consumption without effects on latency to eat nor the motivation to work for food. Consistently, the *in vivo* activity of CeA^Htr2a^ neurons during eating further supports a role for these cells as a circuit node that modulates food consumption, rather than food seeking drive.

The identification of neuronal subpopulations in the CeA that positively regulate food intake has long remained elusive. Recently, a number of genetically-defined CeA neuron populations expressing SOM, CRH, Tac2 and Nts were described to play a role in appetitive behaviors. Nevertheless, the manipulation of these individual neural subpopulations did not affect feeding^15^. Given that these cell populations partially overlap with Htr2a, they likely constitute subsets of the CeA^Htr2a^ population which may explain the lack of effect on eating upon manipulation of the individual populations. Here we could show for the first time that the positive modulation of food intake by the CeA is localized to CeA^Htr2a^ neurons.

Our data support a model where sustained food consumption is promoted by the activity of CeA^Htr2a^ neurons during eating. CeA^Htr2a^ neurons are continually recruited to the active ensemble throughout food consumption and can thereby convey a persistent positive valence signal, which reinforces eating. Given that CeA^Htr2a^ activity increases proximal to eating onset, these neurons are unlikely to reinforce eating by contributing to the postingestive rewarding effects of food consumption associated with nutrient absorption and hormonal changes that occur on timescales of minutes to hours^45,46^. Instead, CeA^Htr2a^ neurons likely promote consumption by influencing food reward during the early phase of consumption and thus may modulate the taste, texture or palatability of the food ^46^. Thus, the positive valence state associated with CeA^Htr2a^ neural activity appears to reinforce eating behaviour after consumption has commenced rather than by driving food seeking itself. This model is consistent with behavioral data showing that activation of CeA^Htr2a^ neurons under conditions where food was non-salient prolonged consumption, while silencing specifically reduced consumption when food was a highly rewarding stimulus. Additionally, the fact that optogenetic stimulation of CeA^Htr2a^ neurons conditioned flavor preference indicates that activity of these neurons enhances the positively rewarding properties of food to reinforce consumption. Thus, CeA^Htr2a^ neurons appear to positively regulate food intake by reinforcement and extension of ongoing food consumption.

A recent study reported a role for GABAergic CeA neurons in pursuit and attack of prey, including artificial prey^47^ without evoking increased food consumption. In contrast, our finding that activation of CeA^Htr2a^ neurons led mice to increase food intake and preferentially engage with and consume a food pellet rather than an inedible item suggests that CeA-mediated consummatory behavior is separable from prey persuit and biting. Importantly, we now reveal that CeA^Htr2a^ neurons mediate consumption and positive reinforcement through its projections to PBN, highlighting that multiple facets of eating behavior are delineated at the level of distinct CeA efferent projections.

Our study outlines how a genetically defined population of CeA neurons is involved in eating-related behavior through both long-range and local circuit mechanisms. Recently, CeA^PKCδ^ neurons were shown to inhibit feeding via local inhibition of other CeA neurons rather than through long-range outputs^11^. Here we show that CeA^Htr2a^ neurons are a source of CeA output that directly modulates consummatory behavior. The functional antagonism of CeA^Htr2a^ neurons and the anorectic CeA^PKCδ^ neurons clearly implicates the CeA in the control of antagonistic consummatory behaviors via reciprocal inhibitory connections between defined CeA cell types. Preferential excitation of one population and thus inhibition of the other, likely serves to flexibly influence consumption, depending on the sensory environment and internal state of the animal.

Although the precise nature of the signals that excite CeA^Htr2a^ neurons are unknown, our data demonstrate that CeA^Htr2a^ neurons are highly innervated by brain regions that process taste and visceral cues^40^, regions associated with energy homeostasis^41^, regions that shows activation upon ingestion of palatable food^42^ as well as regions harboring dopaminergic^44^ and serotonergic neurons^43^. Additionally, we revealed a specific input-output organization of the circuits associated with CeA^Htr2a^ cells. PBN-projecting CeA^Htr2a^ neurons receive a very distinct set of inputs which predominantly target CeA^Htr2a^ neurons compared to CeA^PKCδ^ neurons. How these inputs specifically modulate the activity of CeA^Htr2a^ cells in order to promote eating behaviour through a positive valence mechanism remains an exciting area for future research. Overall, our rabies tracing results highlight that CeA^Htr2a^ neurons are a key node where multiple sensory and internal state modalities are integrated. This complexity in the transmitted signal is reflected in their activity dynamics where CeA^Htr2a^ neurons are recruited during eating but with different latencies, indicating that their activity is not driven by a single input but rather a result of converging, diverse input signals.

A recent study^15^ defined a model in which two circuits from the basolateral amygdala to the CeA may opposingly control appetitive and aversive behaviors. Our data however, indicate that monosynaptic basolateral amygdala inputs represent only a small fraction of total inputs to CeA^Htr2a^ neurons This suggest that appetitive information may converge on the CeA via multiple routes rather than solely through the basolateral nucleus. Together, connectivity mapping supports our major findings by showing that CeA^Htr2a^ cells have all the necessary connections to first assess the valence of food stimuli, and to modulate ongoing eating behavior in return.

Several studies have revealed that the coding of appetitive and aversive behaviors in the CeA is complex as the outcome of neuron activity manipulations of a given subpopulation is dependent on behavioral context^7,9,11,15,48^. In contrast to our finding that CeA^Htr2a^ neurons are active during eating, population activity of these neurons was shown to be reduced upon exposure to innately fearful stimuli^8^. Strikingly, this suggests that CeA^Htr2a^ neurons are implicated in both appetitive and aversive behaviors depending on the behavioral context, highlighting the tremendous behavioral flexibility of CeA neural circuits.

The rewarding nature of CeA^Htr2a^ stimulation suggests that upon induction of feeding, activity of these neurons reinforces eating behavior. This positive valence state may serve as a general reinforcer of consummatory behavior that may be applicable to different salient stimuli encountered in the environment. GABA neurons of the LH have been shown to elicit positive reinforcement and direct consumption toward various stimuli including water and food and to promote interaction with conspecifics^49,50^. This raises the possibility that the CeA may flexibily modulate other reward-related behaviors depending on the environment and internal state of the animal.

In conclusion, we have identified a neural mechanism by which the CeA positively regulates food intake. These findings reveal a new role for the CeA in positive reinforcement of consummatory behavior and identify the underlying neurons and associated circuits. Our findings suggest that the malfunction of these circuits might underlie eating disorders like binge eating. Additionally, our results lay the groundwork for further investigation into how reward processing and behavioral execution is conveyed through interactions between the amygdala and dopaminergic reward system as well as the top-down modulation of gustatory, visceral and taste signals through decending amygdala-brainstem connections. Importantly, our findings also highlight the role of the CeA as a node through which positive and negative valence signals converge and provide an entry point for understanding the interaction between emotional states, food intake and reward.

## Methods

### Animal subjects

The *Htr2a-Cre* BAC transgenic line (STOCK Tg[Htr2a-Cre] KM208Gsat/Mmucd) and *PKCδ-Cre* (Tg(Prkcd-glc-1/CFP,-Cre)EH124Gsat) BAC mice were imported from the Mutant Mouse Regional Resource Center. Td-Tomato (B6.Cg-Gt(ROSA) 26Sortm9(CAG-tdTomato)Hze/J) ^51^ and Rosa26 R ^52^ mouse lines were described previously. Mice were backcrossed onto a C57BL/6 N background.

1–4 month old mice were used in accordance with regulations from the government of Upper Bavaria. Mice used for behavior experiments were singly housed on a 12hr light cycle (7am lights on) cycle with *ad libitum* food access unless food deprived for feeding experiments. All feeding related behavior assays were conducted at a consistent time during light period (8am-1pm). Adult male mice were used for all behavior experiments except for Ca^2+^ imaging experiments where female *Htr2a-Cre* mice were used. Both male and female mice were used for tracing and electrophysiology experiments.

### Viral constructs

The following AAV viruses were produced at the Gene Therapy Center Vector Core at the University of North Carolina Chapel Hill: AAV8-hSyn-DIO-hM3D(Gq)-mCherry, AAV8- hSyn -DIO-mCherry, AAV5-Ef1a-DIO-ChR2-eYFP, AAV5-Ef1a-DIO-eYFP, AAV5-flex-mCherry-dtA, AAV5-ef1a-DIO-npHR3.0-mCherry, AAV1-EF1α-FLEX-TVAmCherry and AAV1-CAG-FLEX-RG. The AAV2/9-Ef1a-DIO-GCaMP6s-eYFP virus was produced at University of Pennsylvania Vector Core. EnvA G-deleted Rabies-GFP used for long range mapping of monosynaptic inputs to CeA^Htr2a^ and CeA^PKCδ^ was produced at the Salk Gene Transfer, Targeting and Therapeutics Core. EnvA G-deleted Rabies-GFP used for TRIO and cTRIO experiments were previously published^35^. AAV8-CAG-Flex^FRT^-G and AAV5-CAG-Flex^FRT^-TC were produced at the Gene Vector and Virus Core of Stanford University School of Medicine. CAV2-Cre and CAV2-Flex^Loxp^-Flp were produced at the Montpellier Vectorology Platform of the UMS Biocampus. HSV-hEF1α-Cre and HSV-hEF1α-LS1L-IRES-flpo were produced at the Viral Gene Transfer Core of the Massachusetts Institute of Technology. AAV-SynMyc ^53^ was a gift from Silvia Arber (FMI, Basel, Switzerland).

### Stereotaxic surgeries

Mice were anesthetized for surgery with isoflourane (1.5–2%) and placed in a stereotaxic frame (Kopf Instruments). Body temperature was maintained using a heating pad. A systemic (Carprofen 5 mg / kg bodyweight) was administered.

Mice for *in vitro* and *in vivo* optogenetic and chemogenetic experiments were bilaterally injected with 0.3µl of virus in the CeA using the following coordinates calculated with respect to bregma: – 1.22mm anteroposterior, ±2.8mm lateral, -4.72mm ventral. In the same surgery, mice for optogenetic experiment were bilaterally implanted with optic fibres (200µm core, 0.22 NA, 1.25mm ferrule (Thor labs)) above the CeA (-4.2mm ventral) or PBN (-5.1mm anteroposterior, ±1.7mm lateral, -3.0mm ventral). Implants were secured with cyanoacrylic glue and the exposed skull was covered with dental acrylic (Paladur). For all other mice, the incision was closed with sutures.

For retrograde tracing experiments 0.15µl retrogradely traveling green or red retrobeads (Lumafluor Inc.) were injected into PBN using the following coordinates from bregma: (-4.8 anteroposterior, ± 1.7 lateral, -3.72 ventral). After 5–7 days post-surgery mice were perfused and brains were processed for histology.

Mice for *in vivo* calcium imaging experiments were injected in the left CeA (coordinates as above) with 0.3µl AAV-GCaMP6s virus. One week later the microendoscope was implanted. Here, a 0.8mm hole was drilled in the skull above the CeA. Debris was removed from the hole and a sterile 20G needle was slowly lowered into the brain to a depth of -4.5 from the cortical surface to clear a path for the lens. The GRIN lens (GLP-0673; diameter: 0.6mm, length: ~7.3mm Inscopix) was slowly lowered into the brain to -4.35 from the cortical surface using a custom lens holder. The lens was secured in place with glue (Loctite 4305) and dental cement (Paladur). A headbar was fixed to the skull adjacent to the lens to assist with mounting of the miniaturized microscope. The exposed top of the lens was protected by a covering of a silicone adhesive (Kwik-cast).

Approximately two weeks after the lens implantation the mice were checked for observable GCaMP6 fluorescence. The mice were headfixed and the top of the lens cleaned of debris. The miniature microscope (Inscopix) with a baseplate (BLP-2, Inscopix) was positioned above the lens such that GCaMP6 fluorescence and neural dynamics were observed. The mice were anesthetized with isoflurane and the baseplate secured with dental cement (Vertise Flow). A baseplate cap (BCP-2, Inscopix) was left in place until imaging experiments.

Mice used to demonstrate monosynaptic inputs to CeA^Htr2a^ and CeA^PKCδ^ were unilaterally or bilaterally injected in the CeA with 0.3–0.4 µL of AAV1-EF1α-FLEX-TVAmCherry and AAV1-CAG-FLEX-RG mixed at a ratio of 1:4 and using the following coordinates calculated with respect to bregma: – 1.22 mm anteroposterior, ±2.8 to 2.9 mm lateral, -4.8 to 4.9 mm ventral. Fourteen days later, 0.3–0.4 µL of EnvA G-deleted rabies-GFP virus was injected into the same area. 7 days after the second injection, the animals were killed and the brains were processed for IHC.

Mice used to demonstrate monosynaptic inputs to PBN-projecting CeA neurons (TRIO experiments) were unilaterally or bilaterally injected in the CeA with 0.3–0.4 µL of AAV1-EF1α-FLEX-TVAmCherry and AAV1-CAG-FLEX-RG mixed at a ratio of 1:4. In the same surgery, they were also injected in the PBN with 0.4 uL of CAV2-Cre and HSV-hEF1α-Cre mixed at a ratio 1:1 and using the following coordinates from bregma: – 5.2 mm anteroposterior, ±1.35 mm lateral, -3.8 to -3.9 mm ventral. Fourteen days later, 0.3–0.4 µL of EnvA G-deleted rabies-GFP virus was injected into CeA. 7 days after the last injection, the animals were killed and the brains were processed for IHC.

Mice used to demonstrate monosynaptic inputs to PBN-projecting CeA^Htr2a^ neurons (cTRIO experiments) were unilaterally or bilaterally injected in the CeA with 0.3–0.4 µL of AAV8-CAG-Flex^FRT^-G and AAV5-CAG-Flex^FRT^-TC mixed at a ratio of 1:1. In the same surgery, they were also injected in the PBN with 0.4 uL of CAV2-Flex^Loxp^-Flp and HSV-hEF1α-LS1L-IRES-flpo mixed at a ratio 1:1. Fourteen days later, 0.3–0.4 µL of EnvA G-deleted rabies-GFP virus was injected into CeA using the same coordinates. 7 days after the last injection, the animals were killed and the brains were processed for IHC.

### Pharmacological treatments

For chemogenetic behavior manipulations, CeA^Htr2a^:: hM3D, and mCherry control mice received an intraperitoneal (IP) injection of CNO (2mg/kg diluted in Saline) or the equivalent volume of saline and allowed to recover in the home cage for 20 minutes prior to the commencement of the experiment. For anorexigenic drug studies, mice were injected IP 20 minutes prior to the experiment: LiCl (150mg/kg) (Sigma), LPS (0.1mg/kg) (Sigma) dissolved in saline or saline control. All drug treatments were delivered in a counterbalanced manner, with three days separating experiments. Behavior experiments were performed with knowledge of the genotype and pharmacological treatment where applicable.

### Optogenetic manipulations

CeA^Htr2a^::ChR2, CeA^Htr2a^::eYFP, CeA^Htr2a^::NpHR, CeA^Htr2a^::mCherry, CeA*^Htr2-Cre+^*::ChR2→ PBN and CeA*^Htr2a-Cre-^*::ChR2→ PBN mice were bilaterally tethered to optic fibre patch cords (Doric Lenses or Thorlabs) connected to a 473nm or 561nm laser (CNI lasers; Cobolt) via a rotary joint (Doric Lenses) and mating sleeve (Thorlabs). For photoactivation experiments, 10ms, 473nm light pulses at 5,10 or 20 Hz and 10-15mW were used. Constant 561nm light at 10mW was used for photoinhibition experiments. The lasers were triggered and pulses controlled by Bonsai data streaming software^54^ and Arduino microcontrollers (www.arduino.cc). For experiments where multiple photostimulation frequencies were tested, the order in which the tests were conducted was randomized.

### Acute brain-slice preparation and electrophysiology

The mice were deeply anesthetized by intraperitoneal injection of Ketamine/Xylazine mixture (100 mg/kg and 10 mg/kg body weight, respectively) and transcardially perfused with ice-cold protective artificial cerebrospinal fluid (aCSF) containing: 92 mM N-methyl-D-glucamine (NMDG), 2.5 mM KCl, 1.25 mM NaH2PO4, 30 mM NaHCO3, 20 mM HEPES, 25 mM glucose, 2 mM thiourea, 5 mM Na-ascorbate, 3 mM Na-pyruvate, 0.5 mM CaCl_2_ 4H_2_O and 10 mM MgSO_4_ 7H_2_O. Coronal brain sections of 250μm thickness were cut with a vibratome (Leica, VT1000S) in ice-cold protective aCSF. For paired recordings thickness was increased to 350μm. Slices were recovered for 15 minutes at 32 °C in regular aCSF containing 126 mM NaCl, 1.6 mM KCl, 1.2 mM NaH2PO4, 1.2 mM MgCl2, 2.4 mM CaCl2, 18 mM NaHCO3, 11 mM glucose, oxygenated with carbogen. After recovery slices were kept at 25 °C until recording.

Slices were visualized using fluorescent microscope equipped with IR-DIC optics (Olympus BX51). All electrophysiological recordings were performed in a chamber constantly superfused with corbogenated regular aCSF at 30–32 °C. Whole-cell voltage, current-clamp or cell-attached recordings were performed with a MultiClamp 700B amplifier and Digidata 1550 (Molecular Devices).

The patch pipette with a resistance of 4–6 MΩ was filled with intracellular recording solution. The intracellular solution for current clamp and paired recordings contained: 130 mM K-Gluconate, 10 mM KCl, 2 mM MgCl2, 10 mM HEPES, 2 mM Na-ATP, 0.2 mM Na2GTP, 0.2% neurobiotin, pH 7.35 and 290 mOsm. The intracellular solution for voltage clamp recordings contained 125 mM CsCl, 5 mM NaCl, 10 mM HEPES, 0.6 mM EGTA, 4 mM Mg-ATP, 0.3 mM Na2GTP, 10 mM lidocaine*N*-ethyl bromide (QX-314), pH7.2 and 290 mOsm. The holding potential for voltage clamp recordings was -70 mV, if not indicated differently. For paired recordings holding potentials ranged from -80 mV to -20 mV. The following drugs were used diluted in aCSF whenever indicated:100 µM Picrotoxin (PTX) and 10 µM NBQX. For confirmation of hM3Dq function in CeAHtr2a neurons, 1 µM of CNO diluted in aCSF was used. Data were sampled at 10 kHz, filtered at 2 kHz and analyzed with pCLAMP10 (Molecular Devices) and Stimfit (http://www.stimfit.org/).

For ChR2-assisted circuit mapping in brain slices a multi-LED array system (CoolLED) connected to the epifluorescence port of the Olympus BX51 microscope was used. 1–2 ms light pulses at *λ =* 470 nm ranging from 1 to 10 mW mm^−2^ was delivered to trigger action potentials in presynaptic cell bodies or axon terminals.

### Behaviour assays

#### Free feeding

For chemogenetic and cell ablation experiments, mice were habituated to the behavior context for daily 10 minute sessions, for two days prior to the experiment. CeA^Htr2a^::hM3D mice were tested in the satiated state at the beginning of the light cycle while CeA^PKCδ^::hM3D and CeA^Htr2a^::dtA mice were tested after 24hr food deprivation. Feeding tests after administration of anorexigenic drugs was conducted after food deprivation. The mice were placed in the behavior box containing two plastic cups in opposing corners, one containing a pre-weighed food pellet. The behavior arena was housed inside a soundproof chamber equipment with houselights and video cameras (TSE Multiconditioning System). For bitter food experiments, food pellets were soaked in 10mM quinine solution (Sigma) for 10 minutes and dried overnight. The mice were allowed to explore the arena for 40 minutes and the remaining food was weighed. The session was video recorded and feeding behavior was scored manually.

For optogenetic experiments, mice were tethered to the optic fibre patch cords and habituated to the context for 15 minutes daily for three days prior to the experiment. On the experiment day the mice were allowed to recover in the behavior context for 5 minutes after tethering. For photostimulation experiments, satiated mice received 20 minutes of photostimulation followed by 20 minutes of no photostimulation. For photoinhibition experiments, mice were food deprived for 24hrs prior to the experiment and received 10 minutes of no photoinhibition followed by 10 minutes of photoinhibition. The quantity of food remaining was measured at the end of each epoch. The session was video recorded and feeding behavior was scored manually.

For experiments comparing consumption of food and clay pellets, clay was prepared by combining kaolin (aluminium silicate hydroxide, Sigma) with 1% gum arabic (Sigma) in distilled water and mixed to form a thick paste. The paste was shaped to the same dimensions as chow food pellets and allowed to dry at room temperature. Three days prior to the experiment, mice were familiarized with the clay pellets in the home cage and habituated to the behavior context for 10 minute daily sessions. The mice were placed in the behavior box containing two plastic cups in opposing corners, each containing a pre-weighed food or clay pellet. After 30 minutes free exploration of the context, the remaining food and clay was weighed

#### Open field

CeA^Htr2a^::dtA, CeA^Htr2a^::hM3D and control mice were allowed to explore a custom plexiglas arena (50cm x 50 cm x 25cm) for 15 minutes. The location of the animal was tracked and the number of entries to the center of the arena (25x25cm square), velocity and distance travelled were assessed using Ethovision XT 11 (Noldus). CeA^Htr2a^::hM3D and CeA^Htr2a^::mCherry mice received an IP injection of CNO (2mg/kg) and allowed to recover in the homecage 20 minutes prior to the experiment.

#### Taste sensitivity

CeAHtr2a::hM3D and mCherry expressing control mice were water deprived overnight prior to the start of the experiment. The mice were trained to drink dH2O from a two-bottle custom licometer (modified from the circuit described in Slotnick., 2009). Each session lasted 1hr per day for five consecutive days. After each session, the mice were allowed *ad libitum* access to water in their homecage for 1hr. On Day 6 mice were injected IP with CNO (2mg/kg) 20 minutes prior to the session where mice where tested for their preference to drink 1mM Quinine (Sigma) solution over dH2O. The preference ratio was calculated by: number of Quinine solution licks/ total number of licks. Licks were timestamped with Arduino microcontrollers and analysed with a custom written Python script.

#### Real-time place preference

CeA^Htr2a^::ChR2, CeA^Htr2a^::eYFP, CeA*^Htr2-Cre+^*::ChR2→ PBN, CeA*^Htr2a-Cre-^*::ChR2→ PBN, CeA^Htr2a^::NpHR and CeA^Htr2a^::mCherry mice were allowed to explore a custom plexiglas two-chambered arena (50cm x 25cm x 25cm). In the case of the photostimulation experiments, mice received 473nm stimulation of 5,10 or 20 Hz in the photostimulated side of the arena which was randomly assigned. For photoinhibition experiments, mice received constant 561nm intracranial light in a randomly assigned compartment. The laser was triggered based on the location of the animal using Bonsai data streaming software and Arduino microcontrollers. The session ran for 20 minutes with the location of the animal, the distance travelled and velocity assessed during the last 15 minutes using Ethovision XT 11 (Noldus).

#### Intracranial self-stimulation

CeA^Htr2a^::ChR2, CeA^Htr2a^::eYFP, CeA*^Htr2-Cre+^*::ChR2→ PBN and CeA*^Htr2-Cre-^*::ChR2→ PBN mice were food restricted overnight prior to the experiment. The assay was conducted over two daily one hour sessions. The mice were placed in a chamber containing a custom two-port nose-poke system modified from https://bitbucket.org/takam/behavioural-hardware. One port was randomly designated the active poke. On Day 1 both active and inactive ports were baited with food treats to encourage exploration. Nose-pokes in the active port resulted in intracranial stimulation (473nm, 10-15mw, 60 x 20 Hz pulse train) while inactive pokes had no consequence. Concurrent with a detected poke, a LED was illuminated below the respective port (1s) and a tone was played (1kHz or 1.5kHz) (1s). Nose-poke time stamps were collected and recorded via Arduino microcontrollers and Bonsai data streaming software and Day 2 data was analyzed using custom written Python script.

#### Progressive ratio 2 (PR2) task

CeA^Htr2a^::hM3D and mCherry mice were food restricted and maintained at 85–90% free-feeding body weight by administering a 2.5g-3.5g food pellet once daily. Mice were trained in daily one hour sessions to nose-poke for food pellets on a fixed ratio 1 schedule (FR1) in the same custom two-port nose-poke system as above. One port was designated the active port. A single nose-poke in the active port which triggered release of one 20mg food pellet (TSE Systems) from a pellet dispenser (Noldus) into a food magazine. Concurrent with a detected poke, a LED was illuminated below the respective port (3s) and a tone was played (1kHz or 1.5kHz) (3s). Nose-poke time stamps were collected and recorded via Arduino microcontrollers and Bonsai data streaming software. Once mice could discriminate between active and inactive pokes by at least 3:1 for three consecutive sessions, mice were trained for three FR5 sessions where five active pokes were required for delivery of a single pellet. This was followed by four sessions on a progressive ratio 2 schedule, where the nose-poke requirement for each successive pellet was increased by two additional responses. Mice were tested for PR2 performance after *ad libitum* access to food following either IP delivery of CNO or saline 20 minutes prior to the session, delivered in a counter-balanced fashion. Breakpoint was considered the highest number of consecutive nose-pokes performed to procure a single food pellet.

#### Conditioned flavor preference

*Ad libitum* fed CeA^Htr2a^::ChR2 mice were habituated overnight to consume two non-nutritive flavoured gels (0.3% grape or cherry sugar-free Kool-Aid (Kraft), 1% Agar (Sigma), 0.15% saccharin (Sigma) in dH2O)). Baseline flavour preference was determined over two consecutive days where mice were tethered to optic-fibre patch cables and habituated for 30 mins and allowed to freely consume both flavours for 15 mins. Baseline preference was the average of the two sessions. Conditioning was conducted over four consecutive days with two-sessions per day. Conditioning session 1: the less-preferred flavour (0.3g) was paired with 25 min intracranial light pulses (473nm, 10-15mW, 20Hz) which commenced 5 min after gel presentation. Conditioning session 2: the mice were presented with the more-preferred (0.3g) flavour for 30 min in the absence of photostimulation. The order of the sessions was inverted each day and occurred 4 hours apart. Conditioned flavour preference was tested the day following the final conditioning session, where the mice were presented with both favours for 15 min. Conditioned preference was averaged from two sessions.

#### Free consumption of palatable reward

CeA^Htr2a^::NpHR mice and controls were food restricted to 85–90% of their free-feeding body weight. Mice were tethered to optic fibre patch cables and allowed to freely consume a palatable liquid reward (Fresubin, 2kcal/ml) from a metal spout for daily 30 minute sessions until stable licking was achieved. The criterion for stable licking was where the number of licks per session over three consective days varied by <± 10% from the first of the three days. Following stable licking acquisition, mice were given *ad libitum* food access and tested for Fresubin consumption during a 20 minute session with constant photoinhibition (561nm, 10mW). Licks were recorded and analysed as for *Taste sensitivity test* experiments.

#### *In vivo* freely moving Ca^2+^ imaging

Two independent groups of CeA^Htr2a^::GCaMP6s mice were used for the free feeding and FR1 imaging experiments. The mice were head fixed and the miniscope was secured in the baseplate holder and the mice were allowed to acclimate in their homecage for 10 minutes prior to the start of imaging. Compressed images were obtained at 20Hz using the Inscopix nVista *HD* V2 software. The LED power was set to 40–60% (0.4–0.6 mW) with the analogue gain set at 1–2.

Mice for the free feeding experiment were acclimated to head-fixation and the weight of the miniscope for 3 daily 15 minute sessions prior to the imaging experiment. For the free-feeding assay, mice were food deprived overnight prior to imaging. The mice were free to explore the arena during the imaging session and consume a food pellet. Mice for the FR1 assay were food restricted to 85–90% of their free-feeding body weight and were trained daily to nose-poke for food rewards on a FR1 schedule with a dummy miniscope in place until they could discriminate between active and inactive pokes by at least 3:1 for three consecutive sessions. For the imaging experiment from three mice: number of active pokes = 12±0, number of inactive pokes = 1.7±0.3.

The average eating bout length for the FR1 experiment was 8.3±2.5s. For both imaging experiments, mouse behavior was recorded using overhead and side-mounted camera and the food contact, eating start and stop events were manually scored from the recorded video. Synchronization of the miniscope software and behavior cameras was achieved using Bonsai data streaming software and a microcontroller (Ardunio).

### Ca^2+^ imaging data analysis

To account for global changes in fluorescence, such as those stemming from neuropil Ca^2+^ signals, time-lapse images of Ca^2+^ activity were filtered using a Fast-Fourier-Transform bandwidth filter from ImageJ (NIH)^33,55^. After motion correction, individual cell filter identification based on combined principal and individual component analysis as well as extraction of raw fluorescence traces was performed using the Mosaic software (v1.1.3; Inscopix)^56^. The Ca^2+^ activity traces and cell masks from each individual unit were visually inspected to ensure that Ca^2+^ signals were obtained from individual neurons. Duplicate or overlapping image filters were removed from the analysis.

*ΔF/F0* was calculated as *(F–F0)/F0*, where *F0* is the lowest 5% of the fluorescence of each Ca^2+^ activity trace ^28^. Normalized *ΔF/F0* was used to transform the range of *ΔF/F0* to [0 1] by the equation:

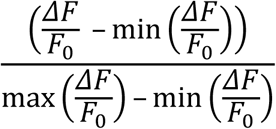

For free-feeding experiments, the first 20 seconds of the first eating bout was compared to the 20 seconds preceding the bout onset. For the classification of neurons all the eating bouts were considered. Average Ca^2+^ fluorescence change of each cell was compared during all the eating bouts longer than 20 seconds to the average fluorescence change of the preceding non-eating bout. For imaging experiments during FR1-task area under the curve was calculated during an event episode and compared to an equal length of baseline before the event onset. For both experiments, eating (F_eating_) and non-eating (F_non-eating_) bout fluorescence changes were compared using Wilcoxon rank-sum test. Cells with significantly different fluorescence changes were categorised as responsive neurons. Next, preference indices (P.I.) for each cell were calculated using the following formula:

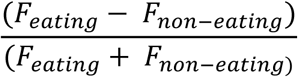

Among the neurons that showed a significant difference in fluorescence changes between non-eating and eating bouts with positive P.I. were categorized as activated during eating, and neurons with negative P.I. were classified as inhibited during eating. All analysis was performed using custom written Matlab and Python scripts. Custom Matlab and Python scripts used for analysis are available upon request.

### Histology

Animals were deeply anesthetized with ketamine/xylazine (100 mg/kg, 16 mg/kg respectively) and transcardially perfused with phosphase-buffered saline (PBS) and then 4% paraformaldehyde (PFA) (w/v) in PBS. Brains were dissected and post-fixed at 4°C in 4% PFA overnight. Brain for cfos immunostaining and in situ hybridization were cryopreserved sequentially in 15% and 30% sucrose in PBS at 4°C before embedding in O.C.T (Fisher Scientific). 50 µm sections were cut with a cryostat (Leica) prior to cfos immunostaing. For in situ hybridization, frozen sections (16μm) were mounted on slides, air dried (30 min at room temperature) and stored at −80°C for later use. All other brains were embedded in agarose after post-fixation and 50 to 100 µm sections were cut with a vibratome (Leica).

### Immunohistochemistry

For immunostaining, sections were washed in 1X PBS 0.5% TritonX-100 and blocked at room temperature for two hours in 1% bovine serum albumin (BSA) diluted in 1X PBS 0.1% TritonX-100 and then incubated in primary antibody at 4°C overnight. Primary antibodies: rabbit anti-cfos (1:1000) (Santa Cruz); mouse anti-PKCδ (1:100) (610398, BD Biosciences), chicken anti-LacZ (1:200) (ab9361, Abcam), rabbit anti-SOM (1:1000) (T-4103, Peninsula Laboratories International), goat anti-CGRP (1:500) (Abcam, ab36001), rabbit anti-myc (ab9106, Abcam). Following 3x 15 minute 1X PBS 0.1% TritonX-100 washes, sections were incubated in secondary antibody at for 2 hours at room temperature. Secondary antibodies: Alexa Fluor donkey anti-rabbit/mouse/goat 488/Cy3/647, (1:500) (Jackson). After 3x15 minutes 1X PBS 0.1% TritonX-100 washes, sections were incubated in DAPI and coverslipped (Dako).

For detection of cfos in CeA^Htr2a^:: hM3D and mcherry control mice CNO (2mg/kg) was injected IP and animal were perfused two hours later. For detection of cfos in CeA^Htr2a^:: ChR2 and eYFP control mice, animals received 20 minutes 20Hz, 10mW photostimulation and were perfused one hour later.

For recovery of neurobiotin filled neurons after whole cell recordings, acute brain slices were fixed in 4% PFA at room temperature for 30–45 minutes. Fixed slices were kept in 0.1M PB (80mM Na2HPO4, 20mM NaH2PO4) until being processed for immunohistochemistry as described above. Slices were then washed in 0.1 M PB and incubated for 48 hours at 4 °C with mouse anti-PKCδ (1:100) (610398, BD Biosciences) or goat anti-CGRP (1:500) (Abcam, ab36001) diluted in 0.05 M TBS (42 Trizma HCl mM, 8 mM Trizma base and 154mM NaCl) with 0.5% Triton X-100 added. Afterwards, slices were washed in 0.1M TBS and incubated overnight in secondary antibodies (1:500) (Jackson) and fluorophore conjugated streptavidin (1:2000) (Jackson) diluted in 0.05 M TBS with 0.5% Triton X-100. The next day, slices were washed in 0.1M PB and mounted using RapiClear (SunJin Lab Co) and imaged the day after.

### *In situ* hybridization

Two colour FISH was performed on fixed frozen sections from *Htr2a-cre;LacZ, Htr2a-cre;tdTomato* mice or *PKCδ-Cre* mice that underwent stereotaxic surgeries to demonstrate monosynaptic inputs to CeA^PKCδ^ using the proprietary probes and methods of Advanced Cell Diagnostics (Hayward, CA, United States) (ACD Technical notes #320535 for sample preparation, and #320293 for Multiplex fluorescent labeling, http://www.acdbio.com/technical-support/downloads).

Briefly, OCT was removed with PBS before pretreatment with ACD proprietary reagents Pretreat 2 and Pretreat 4. Sections were boiled for 5 min in Pretreat 2 buffer, washed in distilled water and 100% ethanol. Sections were air dried before incubation with Pretreat 4 for 30 min at 40°C in a HybEZ humidified incubator (ACD). We performed single or dual probe labeling, using probes for Htr2a (Mm-Htr2a-C1, #401291), LacZ (Ecoli-lacZ-C3, #313451-C3), TAC2 (Mm-Tac2-C2, #446391) and CRH (Mm-CRH-C2, #316091). C1 probe was ready to use. When used in combination with C1 probe, C2 and C3 probes were diluted 50 times in C1 probe. When used alone, C2 and C3 probes were diluted 50 times in the probe diluent buffer. Tissue sections were incubated in probe mix for 2 hr at 40°C in the HybEZ humidified incubator. Sections were washed in ACD Wash Buffer (2 × 2′) then sequentially incubated in ACD proprietary reagents AMP1-FL (30 min), AMP2-FL (15 min), AMP3-FL (30 min) and Amp 4 Alt B-FL AMP4-FL (15 min) with two washes (2 min) between each step. Brain sections from *Htr2a-cre;LacZ* mice were then labeled with DAPI and coverslipped using Fluorescent Mounting Medium (Dako). Brain sections from tdTomato mice or PKCδ-Cre mice that underwent stereotaxic surgeries to demonstrate monosynaptic inputs to CeA^PKCδ^ were blocked 2 hr at RT with 0.2% BSA and 5% donkey serum. Sections were then incubated with a mouse anti-GFP (1:500) (632381, Clontech) and, or a rabbit anti-mcherry (1:500) (ab 167453, Abcam) in 0.1% TritonX-100 and 0.2% BSA at 4°C overnight. Sections were washed 3 times 15 minutes in PBS and incubated with secondary antibodies: Alexa Fluor donkey anti-rabbit/mouse 488/Cy3/647, (1:300) (Jackson) for 2 hours at room temperature. Following 3 times 15 minutes washes, sections were incubated in DAPI and coverslipped (Dako).

### Microscopy

Fluorescent Z-stack images were acquired using a Leica SP8 confocal microscope and a 20X/0.75 IMM objective (Leica). Epifluorescent images were obtained using an upright epifluorescent microscope (Zeiss) using 5X/0.15 or 10X/0.3 objectives (Zeiss). Images were minimally processed using ImageJ software (NIH) to enhance brightness and contrast. Median and mean filters were used to reduce noise. For colocalization analyses of *Htr2a-Cre* neurons with PKCδ and SOM, retrobead tracing experiments and for detection of cfos, sections were quantified from anterior to posterior CeA (bregma -1.22 to -1.58 mm) (n = 3 sections from 3 mice) using ImageJ. For quantification of *Htr2a-Cre::tdTomato* neurons that remained in *Htr2a-Cre::dtA* mice, mCherry (infected but unablated) expressing neurons were distinguished from tdTomato neurons using an upright epifluorescent microscope (Zeiss) (n = 3 sections from 3 mice). For quantification of local monosynaptic inputs to CeA^Htr2a^ and CeA^PKCδ^ neurons and PBN-projecting CeA neurons, sections were quantified from anterior to posterior (bregma -0.94 to -1.82 mm) (n = 3–4 50 µm sections per animal and for each marker) using ImageJ. For colocalization analyses of β-Gal reporter with Htr2a mRNA and tdTomato reporter with TAC2 and CRH mRNA, sections were quantified from anterior to posterior (bregma -1.22 to -1.58 mm) (n =3 12 µm sections per animal) using ImageJ. For quantification of starter cells in the CeA, every section or every second section from -0.94 to -1.82 mm posterior to bregma was quantified using ImageJ.

### Long-range mapping of monosynaptic inputs

6 CeA^Htr2a^, 5 CeA^PKCδ^, 5 PBN-projecting CeA and 4 PBN-projecting CeA^Htr2a^ traced neurons brains were chosen based on high tracing efficiency and large number of starter cells mainly restricted to the CeA.

For quantifications of subregions, boundaries were based on the Allen Institute’s reference atlas ^57^ with consultation of Paxinos and Franklin (2013) atlas ^58^. Our definition of VP, anterior and posterior PVT, SNL and DR, was exclusively according to Paxinos and Franklin (2013) atlas.

For quantifications within all subregions, every section was quantified but only the input neurons found ipsilateral to the injection site were counted. Input regions that were part of the amygdala complex except for the LA, BLA and BLP were excluded from the analysis, namely the basomedial amygdala nucleus, intercalated amygdala nucleus, central amygdala nucleus, medial amygdala nucleus, cortical amygdala area, posterior amygdala nucleus, piriform-amygdala area, postpiriform transition nucleus. Moreover, a small number of starter neurons were found in neighboring nuclei: the globus pallidus (GP), caudoputamen (CP), and the very lateral part of the lateral hypothalamus. Although these neurons accounted for a small portion of the total starter neurons, we excluded input cells from these areas for the analysis.

The numbers of input neurons for each experiment were normalized to the total number of inputs in each animal. Areas that contained <1% of the total inputs in both genotypes were excluded. One-way ANOVA with Bonferroni post-hoc test was used to compare within the Htr2a and PKCδ groups whether the proportion of inputs coming from each region is statistically different to their respective average proportions of all input regions. Average proportions of all input regions to CeA^Htr2a^ or CeA^PKCδ^ were calculated by considering only regions that give <1% of the total inputs to CeA^Htr2a^ or CeA^PKCδ^ respectively. Coefficients of variation for each region giving more than 1% of the total inputs to CeA^Htr2a^ and, or CeA^PKCδ^ neurons were calculated by dividing the standard deviations by the means. All graphs and analysis described above were performed using Prism software (GraphPad).

Pairwise correlations (Pearson coefficient) as well as p values from a unpaired two-tailed t-test were calculated in Excel (Microsoft) using unormalized data. Hierarchically clustered heatmaps and dendrograms representing high correlation or anticorrelation between regions were created using R (http://www.r-project.org/). Linear regressions were performed using Prism software (GraphPad).

### 3D-reconstruction of CeA^Htr2a^ axonal projections

The protocol was adapted from a previously described protocol ^59^. *Htr2a-Cre* mice stereotaxically injected into the CeA with an AAV-mcherry virus, were transcardially perfused with 20mL of ice cold 1X PBS followed by 20mL of an ice cold hydrogel monomer solution based on 2% acrylamide (161–0140, Bio-Rad), 0.025% Bisacrylamide (1610142, BioRad), 4% PFA, 0.25% VA-044 initiator (w/v) (27776–21-2, Wako) in 1X PBS at a speed of 4mL/minute. After perfusion, brains were incubated for 3 days at 4°C. Samples were then polymerized for 3 hours at 37°C. The embedded samples were extracted from the gel and cut using a vibratome in 2mm sections. 2mm sections were incubated with the clearing solution (4% SDS, 200mM Boric Acid, pH 8.5) until the sections became cleared (about 2 weeks). Sections were then washed for at least 3 days in 1X PBS, 0.1% TritonX-100 at 37°C in order to remove residual SDS. Sections were finally incubated in a refractive index matching solution (RapiClear, RI=1.47, SunJin Lab Co) for 8 hours (up to 1 day) at room temperature. Samples were finally mounted in fresh RapiClear between two coverslips separated by iSpacers (SunJin Lab Co) and imaged using a Leica TCS SP8 microscope (Leica Microsystems) with a 10X/0.30 objective.

Processing of the 3D images was done using the Amira software (Visage Imaging Inc). Serial stack images were first roughly registered with respect to each other and transformed into one coordinate system via the Transform Editor. Serial stack images were then concatenated and aligned in 2D using the Align Slices module (all transformations were rigid). Sigmoid intensity remapping was applied in order to specifically raise the intensity range of the axonal fibers over the background. Manual segmentation of CeA^Htr2a^ axonal projections and of the edge of the sections was done using the Segmentation Editor. 3D rendering of the brain surface was generated using the Surface view module. 3D impression of CeA^Htr2a^ axonal projections as well as colour coding of the intensity of fluorescent pixels was completed using the Volume Rendering module.

### Statistics

No statistical methods were used to predetermine sample size. The number of samples in each group were based on published studies. Behavior experiments were conducted by an investigator with knowledge of the animal genotype and treatment. The need for post-hoc verification of virus expression and optic fibre placement ensured data collection was unbiased. For most behavior experiments, physiology and *in vivo* imaging, custom written Python scripts, behavioral tracking software and automated behavioral analysis was used to obtain and analyse the data in an automated and unbiased manner. For behavior experiments, littermate animals were randomly assigned to the experimental group and were identified by unique identification number. Following the conclusion of experiments with animals, virus expression and optic fibre and GRIN lens placement was verified. Mice in which either the virus expression or optic fibre was not appropriately located were excluded from analysis. Data presented as box–whisker plots display median, interquartile range and 5th–95th percentiles of the distribution. Data presented as bar and line graphs indicate mean ± SEM. Pairwise comparisons were calculated by un-paired or paired two-tailed t-tests and multiple group data comparisons were calculated by one-way or two-way ANOVA with Bonferroni post-hoc test. Normality was assessed using Shapiro-Wilk tests. In the case where normality tests failed, Mann-Whitney or Wilcoxon rank-sum tests were used. Statistical analysis were performed using Graphpad Prism 6.0, Matlab or Python. Significance levels are indicated as follows: *p<0.05, **p<0.01, ***p<0.001, ****p<0.000. See Supplemetary Statistics Table.

## Data and code availability

All relevant data and custom-written analysis code are available from the corresponding author upon reasonable request.

## Author Contributions

A.M.D, H.K, M.P, M.M, J.G and C.S designed and analyzed experiments. A.M.D performed behavior experiments. H.K performed electrophysiology and assisted with behavior hardware design. M.P performed rabies tracing. H.K and M.P performed other tracing experiments. C.S performed paired-recordings and hM3D *ex vivo* verification. A.M.D, H.K, M.P and P.L.A.M performed histology. A.M.D, H.K, M.M and J.G performed Ca^2+^ imaging experiments. K.K.C. provided the rabies virus and expertise for its use for monosynaptic tracing. R.K and A.L supervised experiments. A.M.D, H.K, M.P and R.K wrote the manuscript with input from all other authors.

## Acknowledgments

We thank Tommaso Caudullo, Elke Fejzulahi and Bettina Hoisl for their help with the management of the animal colony, Robert Kasper for technical help with the *in vivo* optogenetics setup, Tobias Ruff and Daniel del Toro for help with tissue clearing, Martin Schwarz (Max Planck Institute for Medical Research, Heidelberg) for providing Cre-dependent AAV-mCherry virus, Silvia Arber (Friedrich Miescher Institute for Biomedical Research, Basel) for providing Cre-dependent AAV- synaptophysin-myc virus, and Alex Ghanem for initial help with rabies virues. We thank Pankaj Sah and Nadine Gogolla for critical reading of this manuscript. Modified rabies virus were provided by the GT3 Core Facility of the Salk Institute with funding from NIH-NCI CCSG: P30 014195, an NINDS R24 Core Grant and funding from NEI. This study was supported by the Max-Planck Society and the Deutsche Forschungsgemeinschaft (Synergy) (to R.K.).

